# Efficient Game-Theoretic Explanations for Tree-Based Ensembles via Owen Values

**DOI:** 10.64898/2026.08.12.744440

**Authors:** Hyunwook Koh

**Affiliations:** Department of Applied Mathematics and Statistics, The State University of New York, Korea, Incheon, South Korea

**Keywords:** Owen Value, Explainable Artificial Intelligence, Game-Theoretic Feature Attribution, Local Explanation, Feature Importance, Tree-Based Ensembles

## Abstract

Shapley-value-based explanations, notably SHAP (SHapley Additive exPlanations), have gained prominence as a principled game-theoretic framework for local explanations and global feature importance. While exact Shapley value computation is exponential in feature count, TreeExplainer exploits the recursive structure of decision trees to achieve polynomial-time computation for tree-based ensembles. In many scientific applications, however, features are naturally organized into *a priori* groups reflecting domain knowledge, requiring explanations both across and within groups. The Owen value extends the Shapley value through a two-stage allocation rule that incorporates group structure while preserving fairness properties; yet, efficient algorithms for its computation remain limited. In this paper, we propose exact and Monte Carlo algorithms for computing Owen values in tree-based ensembles by combining hierarchy-guided group aggregation with tree-aware dynamic programming. The exact algorithm computes Owen values without sampling under the path-dependent characteristic function, which approximates the conditional expectation, whereas the Monte Carlo algorithm provides a scalable approximation that is unbiased for any prespecified sampling budget and converges almost surely as the sampling budget increases. We also provide global importance measures and visualization tools for structured, multi-resolution explanations. The proposed algorithms and tools are collectively referred to as TreeOwen. Through simulation experiments, we demonstrate the numerical accuracy and substantial computational gains of TreeOwen. We illustrate its practical utility using immunotherapy metagenomic data, showing how microbial genera (groups) and species (features) contribute to patient recovery.

## 1 Introduction

In modern machine learning, Shapley-value-based explanations have been introduced by Štrumbelj and Kononenko (2010, 2014) and subsequently unified and popularized through SHAP (SHapley Additive exPlanations) (Lundberg and Lee, 2017), which enables both local explanations at the individual observation level and global measures of feature importance. SHAP has since become a central tool in explainable artificial intelligence (XAI), particularly for complex nonlinear models where interpretability is otherwise limited. A key advance came with TreeExplainer (Lundberg et al., 2020), which exploits the recursive structure of decision trees via dynamic programming to reduce exact Shapley value computation from exponential to polynomial time for tree-based ensembles, such as gradient boosting machines (Friedman, 2001) and random forests (Breiman, 2001), making Shapley-based feature attribution no longer the primary computational bottleneck for this class of models.

In many scientific applications, however, features are naturally organized into *a priori* groups reflecting domain knowledge. Examples include microbial taxa in metagenomics, pathway-level gene sets in genomics, company- or industry-level factors in financial markets, and word- or sentence-level representations in natural language processing. In such settings, interpretability is required not only at the individual feature level but also at the group level, and ideally across multiple resolutions simultaneously: local attributions for individual observations and global importance measures summarized across all observations.

The Owen value (Owen, 1977) extends the Shapley value to games with *a priori* group structure via a two-stage allocation rule: the total value is first distributed across groups and then allocated among individual features within each group. This extension preserves the fairness axioms of the Shapley value while explicitly encoding the domain-imposed group structure, thereby providing a theoretically grounded framework for structured, multi-resolution attribution that ensures coherent explanations at both the group and feature levels. Despite its theoretical appeal, scalable algorithms for computing Owen values in modern machine learning models remain limited. Exact Owen value computation requires enumerating all outer group subsets as well as all inner feature subsets within each group, leading to complexity that grows exponentially in both the number of groups and the within-group sizes. While TreeExplainer resolved this problem for standard Shapley values in tree-based ensembles (Lundberg et al., 2020), analogous algorithmic developments for Owen values in such models remain underdeveloped, limiting the routine adoption of structured interpretation in practice.

In this paper, we address this limitation by developing computationally efficient algorithms for Owen values in tree-based ensembles, collectively referred to as TreeOwen. In this framework, the grouping is supplied *a priori* by domain knowledge to define the explanation question rather than the prediction rule, and is not assumed to be represented internally by the trained model. We propose an exact algorithm that decomposes the computation into hierarchy-guided aggregation across groups and tree-aware dynamic programming within each outer group context, thereby avoiding explicit outer group subset enumeration and substantially reducing the combinatorial burden while preserving exactness with respect to the path-dependent characteristic function (Lundberg et al., 2020), which approximates the conditional expectation (Štrumbelj and Kononenko, 2010, 2014; Lundberg and Lee, 2017). To further improve scalability when within-group sizes are large, we also propose a Monte Carlo algorithm that replaces exhaustive inner enumeration with a permutation-based estimator, which is unbiased for any prespecified sampling budget and converges almost surely as the sampling budget increases. In addition to the computational framework, TreeOwen provides global importance measures and feature- and group-level visualization tools to support structured, multi-resolution interpretation of tree-based ensembles. Through simulation experiments, TreeOwen is evaluated along two dimensions: numerical accuracy and computational efficiency, relative to enumeration baselines. The practical utility of TreeOwen is further illustrated through an application to metagenomic data from a cancer immunotherapy study (Limeta et al., 2020), demonstrating how microbial genera (groups) and species (features) contribute to patient recovery.

The grouped attribution computed by TreeOwen can be related to, and distinguished from, two strands of prior work. The first is the summed Shapley attribution, a general strategy that simply sums feature-level Shapley values over each group. It can be combined with any per-feature Shapley estimator, including KernelSHAP (Lundberg and Lee, 2017), TreeSHAP (Lundberg et al., 2020), FastSHAP (Jethani et al., 2022), and conditional KernelSHAP (Aas et al., 2021). These estimators differ in the characteristic functions they evaluate, including conditional, marginal/interventional, and tree-path-dependent formulations. Among these alternatives, TreeOwen builds on the path-dependent recursion of TreeSHAP (Lundberg et al., 2020), which incorporates dependence information encoded in the split structure and node cover statistics of the fitted tree without requiring explicit estimation of a joint or conditional feature distribution, and supports the repeated within-context evaluations required by our algorithms. Although the summed Shapley attribution is simple, widely used, and broadly applicable, it lacks an axiomatic characterization as a grouped solution concept. In contrast, the Owen value that TreeOwen computes is axiomatically characterized for games with *a priori* unions (Owen, 1977; Alonso-Meijide et al., 2009). The second strand, known as groupShapley (Jullum et al., 2021), instead computes Shapley values with groups as players, yielding group-level attributions but no within-group decomposition. In contrast, the Owen value provides both resolutions: its group-level totals coincide with such group-level Shapley values, while it additionally allocates each group’s contribution among its member features.

The remainder of the paper is organized as follows. Section 2 introduces the cooperative-game formulation and develops the TreeOwen algorithms and their theoretical properties. Section 3 presents simulation studies and the real data application. Section 4 summarizes the main findings, discusses limitations and directions for future research, and concludes the paper.

## 2 Methods

### 2.1 Cooperative Game, Shapley Value, and Owen Value

#### Cooperative Game for Tree-Based Ensembles

Let *p* ∈ ℕ denote the number of input features and define the feature index set *N* := {1, …, *p*}. Let *f* : ℝ^*p*^ → ℝ be a trained tree-based ensemble, such as a gradient boosting machine (Friedman, 2001) or a random forest (Breiman, 2001). The predictor is written in additive form as

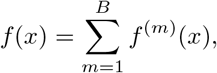

where each *f* ^(*m*)^ denotes an individual decision tree (Breiman et al., 1984) and *B* is the number of trees; where applicable, boosting weights and learning-rate factors are absorbed into each *f* ^(*m*)^.

For a target observation *x* ∈ ℝ^*p*^, the local explanation problem is formulated as a cooperative game (*N, v*_*x*_), where the characteristic function *v*_*x*_ : 2^*N*^ → ℝ assigns a value to each subset of features. Following the standard conditional expectation semantics (Štrumbelj and Kononenko, 2010; Štrumbelj and Kononenko, 2014; Lundberg and Lee, 2017), the characteristic function is defined as

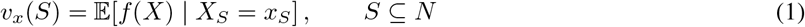

TreeOwen uses the path-dependent characteristic function of TreeSHAP (Lundberg et al., 2020) (Algorithm 1) as an approximation to the conditional-expectation target in Eq. (1). For this path-dependent game, *v*_*x*_(*N*) = *f* (*x*), while *v*_*x*_(∅) is the baseline induced by the fitted tree structure and node cover ratios (Lundberg et al., 2020). Alternatively, the marginal/interventional characteristic function (Lundberg and Lee, 2017; Janzing et al., 2020) or a conditional characteristic function (Aas et al., 2021) may be substituted for *v*_*x*_; however, none of these is guaranteed to recover the conditional expectation in Eq. (1) exactly in practice, because the true conditional distribution is unknown. Under the additive decomposition of *f*, the characteristic function inherits linearity 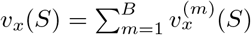, which implies that all solution concepts introduced below decompose additively across trees.

#### Shapley Value and Owen Value

For *i* ∈ *N*, the Shapley value (Shapley, 1953) is given by

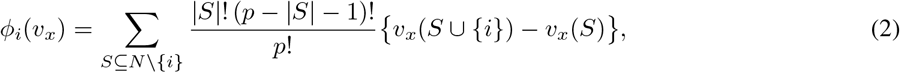

which is the unique solution satisfying standard axioms (efficiency, symmetry, additivity, and the null-player property).

Now suppose that the features are partitioned into *K* disjoint groups *G* = {*G*_1_, …, *G*_*K*_} with 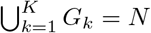, and define the group index set *K* := {1, …, *K*}. This induces a group-level game 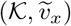 with

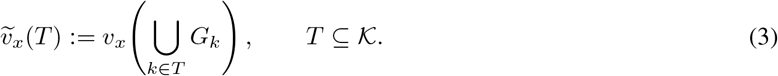

For a feature *i* ∈ *G*_*k*_, the Owen value (Owen, 1977) is given by

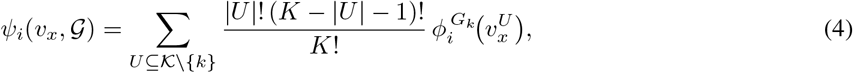

where 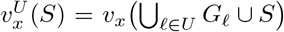 for *S* ⊆ *G*_*k*_; equivalently, setting *C* := ∪_*ℓ*∈*U*_ *G*_*ℓ*_, we write 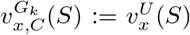, and 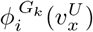 denotes the Shapley value of player *i* in the |*G*_*k*_|-player cooperative game 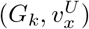, given explicitly by

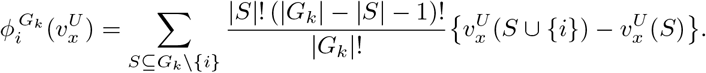

This allocation is the unique solution satisfying the following five axioms (Owen, 1977):

i. Group-level efficiency: 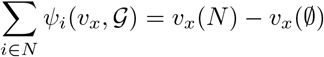;
ii. Feature-level efficiency: 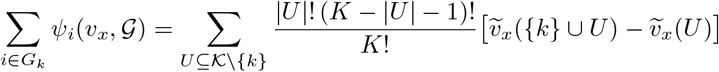 for each *G*_*k*_; that is, the within-group sum equals the Shapley value of group *k* in the group-level game 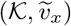;
iii. Symmetry: if 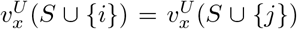 for all *S* ⊆ *G*_*k*_ \ {*i, j*} and all *U* ⊆ *K \* {*k*}, then *ψ*_*i*_(*v*_*x*_, *G*) = *ψ*_*j*_(*v*_*x*_, *G*);
iv. Additivity: *ψ*_*i*_(*v*_*x*_ + *w*_*x*_, *G*) = *ψ*_*i*_(*v*_*x*_, *G*) + *ψ*_*i*_(*w*_*x*_, *G*);
v. Null-player property: if 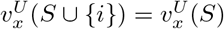 for all *S* ⊆ *G*_*k*_ \ {*i*} and all *U* ⊆ *K \* {*k*}, then *ψ*_*i*_(*v*_*x*_, *G*) = 0.

The Owen value yields a two-resolution attribution. At the outer resolution, the group total 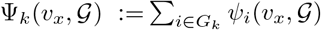 quantifies the contribution of group *G*_*k*_ as a single unit, with each group entering or leaving an outer coalition atomically. At the inner resolution, the within-group values 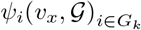 distribute Ψ_*k*_ among the individual features of *G*_*k*_. Features are thus treated jointly only at the outer level, while their individual roles are resolved at the inner level. We refer to this linked allocation at the group and feature levels as a multi-resolution explanation.

How the Owen value relates to the ordinary Shapley value—and to its group-wise sum, the summed Shapley attribution 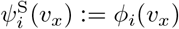—depends on the group structure and the interactions in *v*_*x*_. First, when every group is a singleton, the Owen value reduces to the Shapley value, *ψ*_*i*_ = *ϕ*_*i*_ (Owen, 1977). Second, more generally, when *v*_*x*_ is additive across groups so that no interaction crosses a group boundary, the inner allocation is context-independent and the Owen value coincides with the summed Shapley attribution, giving 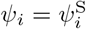 feature by feature. Finally, when interactions cross group boundaries, the Owen value in general no longer coincides with the summed Shapley attribution, and the two are best understood not as competing estimates of a single quantity but as distinct solution concepts that satisfy different axioms.

##### Proposition 1

(Owen versus summed Shapley).

*The summed Shapley attribution* 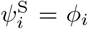 *satisfies (i) group-level efficiency and (iv) additivity, but does not in general satisfy (ii) feature-level efficiency, (iii) symmetry, or (v) the null-player property*.

Properties (i) and (iv) follow from the efficiency and additivity of the Shapley value; the other three are stated relative to the group structure—through the quotient game 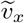 in (ii), and over group-complete outer contexts 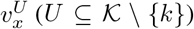 in (iii) and (v)—whereas the Shapley value also responds to partial cross-group coalitions, so the summed Shapley attribution can violate (ii), (iii), and (v) when such interactions are present. The proof, with explicit counterexamples, is given in the *Supplementary Materials*. The summed Shapley attribution and the Owen value thus answer an ungrouped and a group-structured attribution question, respectively, and are complementary. Because the summed Shapley attribution and the Owen value define different attribution targets, we treat the former as a conceptual comparator rather than a numerical accuracy reference; exact Owen enumeration is used to validate the numerical accuracy of TreeOwen.

### 2.2 Naive Enumeration Methods

Two naive enumeration approaches serve as exact reference implementations for the Owen value.

i. *Model-agnostic enumeration*. This approach treats the ensemble as a black box and exhaustively enumerates all outer group subsets *T* ⊆ *K \* {*k*} and inner feature subsets *S* ⊆ *G*_*k*_ \ {*i*}. The overall complexity is 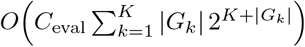, with *O*(*p*) working memory.
ii. *Tree-aware enumeration*. This approach performs the same enumeration but replaces black-box evaluation with tree-aware dynamic programming (Section 2.3), reducing per-evaluation cost to *O*(*M*) and overall complexity to 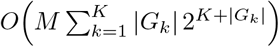, with *O*(*M*) working memory.

Both methods are exponential in *K* and |*G*_*k*_| and are infeasible at scale.

### 2.3 TreeOwen: Exact Algorithm

#### Step 1: Hierarchy-Guided Aggregation across Groups

Direct evaluation of the Owen value in Eq. (4) requires enumerating all 2^*K*−1^ outer group subsets *U* ⊆ *K \* {*k*}, which grows exponentially in *K*. TreeOwen avoids this by introducing an auxiliary balanced binary tree ℬ over the *K* groups to organize the outer Shapley-weighted summation.

For any node *u* in ℬ, let ℒ (*u*) ⊆ *K* denote the set of group indices at the leaves of *u*, and define 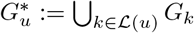. For an internal node *u* with children (*u*_*L*_, *u*_*R*_), the induced sub-partition over 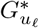 is {*G*_*k*_ : *k* ∈ ℒ (*u*_*ℓ*_)} for each *ℓ* ∈ {*L, R*}. For any leaf *G*_*k*_, let *d*_*k*_ denote its depth in ℬ.

The outer aggregation proceeds top-down over ℬ, maintaining a context *C* disjoint from 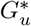. At each internal node, the sibling subtree is either excluded or included in *C*, each with weight 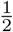. This yields 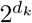 contexts for group *G*_*k*_. For a balanced ℬ, *d*_*k*_ ≤ ⌈log_2_ *K*⌉, giving at most 2*K* − 1 contexts per leaf instead of 2^*K*−1^. We use a balanced ℬ because it minimizes the maximum leaf depth and thereby bounds the number of contexts visited per leaf (Lemma 1); this is a computational choice for controlling cost rather than a claim of unique optimality. Upon reaching leaf *G*_*k*_, the within-group solver of Step 2 is invoked under context *C*, and results are aggregated with the corresponding weights. The procedure is given in Algorithm 2.

#### Step 2: Tree-Aware Dynamic Programming and Within-Group Enumeration

Characteristic function values *v*_*x*_(*S*) in Eq. (1) are evaluated via tree-aware dynamic programming, rather than treating the ensemble as a black box (Lundberg and Lee, 2017; Lundberg et al., 2020). The evaluator is given in Algorithm 1.

i. *Per-tree recursion*. Consider a decision tree with node set *V*. Each node *u* ∈ *V* is either an internal node that splits on feature *j*(*u*) with threshold *τ* (*u*) and children *c*(*u*), or a leaf with value val(*u*). For *S* ⊆ *N*, let *V*_*u*_(*S*) denote the path-dependent value of the subtree rooted at *u*. Then

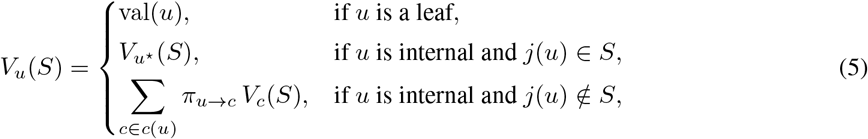

where *u*^⋆^ is the child consistent with *x*_*j*(*u*)_, and *π*_*u*→*c*_ = *n*_*c*_*/n*_*u*_ (or 1*/*| *c*(*u*)| when *n*_*u*_ = 0). This recursion is implemented as NodeDP; EvalV sums over all *B* trees. Correctness under degenerate cover and missing values is given in Proposition 2 in *Supplementary Materials*.
ii. *Handling missing features*. A feature is *known* if it belongs to *S*. If *j* ∈ *S* but *x*_*j*_ is missing, the default missing direction is used when available; otherwise *j* is treated as unknown.
iii. *Extension to ensembles*. By additivity, EvalV sums NodeDP over all trees, and the per-evaluation cost is *O*(*M*).
iv. *Memoization*. For a fixed context *C*, TreeOwen evaluates *v*_*x*_(*C* ∪ *S*) for all *S* ⊆ *G*_*k*_. A feature–tree inverted index identifies trees affected by each feature, so when extending *S* to *S* ∪ {*i*}, only trees splitting on *i* are recomputed, while others are cached.
v. *Exact within-group Shapley enumeration*. For *G*_*k*_ and context *C* ⊆ *N* \ *G*_*k*_, define 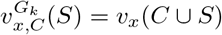. For *i* ∈ *G*_*k*_,

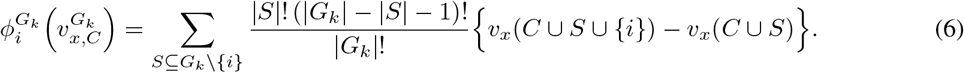

All 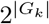 subsets are enumerated, with *v*_*x*_(*C* ∪ *S*) evaluated via EvalV and memoization. The total cost is 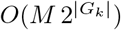. This within-group computation is exact under the given context *C*; its cost, however, grows exponentially in |*G*_*k*_|, which is precisely the regime handled by the Monte Carlo algorithm of Section 2.4.

#### Theoretical Properties of the Exact Algorithm

Correctness and runtime guarantees for Algorithm 2 are established below. Proofs of all results are provided in *Supplementary Materials*.

##### Lemma 1

(Context Count and Weight Partition).

*Let* ℬ *be a balanced binary tree with K* ≥ 1 *leaves, and let d*_*k*_ *denote the depth of the leaf corresponding to group G*_*k*_. *For any leaf G*_*k*_:

i. *(Count)* Recurse *visits G*_*k*_ *under exactly* 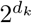 *distinct outer contexts. Hence* 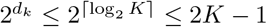.
ii. *(Weight partition) If* 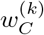 *denotes the weight assigned to context C when G*_*k*_ *is visited, then* 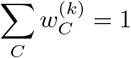.

##### Theorem 1

(Correctness).

*Let G* = {*G*_1_, …, *G*_*K*_} *be a partition of N and let v*_*x*_ : 2^*N*^ → ℝ *be any characteristic function. For every i* ∈ *G*_*k*_, *Algorithm 2 returns* 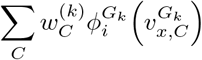, *where the contexts and weights are as specified in Lemma 1*. *The proof is given in Supplementary Materials*.

By Theorem 1 and the efficiency axiom of the Owen value (Owen, 1977), the algorithm output satisfies ∑_*i* ∈*N*_ *ψ*_*i*_(*v*_*x*_,*G*) = *v*_*x*_(*N*) − *v*_*x*_(∅) = *f* (*x*) − *v*_*x*_(∅), where *v*_*x*_(∅) is the baseline induced by the path-dependent characteristic function.

##### Theorem 2

(Runtime Complexity).

*Under the conditions of Theorem 1, with* ℬ *a balanced binary tree and characteristic function values evaluated via* EvalV *(Algorithm 1), the total number of* EvalV *calls is at most* 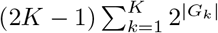, *and the total runtime is*

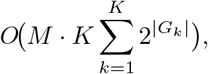

*where M is the total node count across all B trees*.

##### Algorithm 1

Path-Dependent Characteristic Function via Tree DP (NodeDP / EvalV)

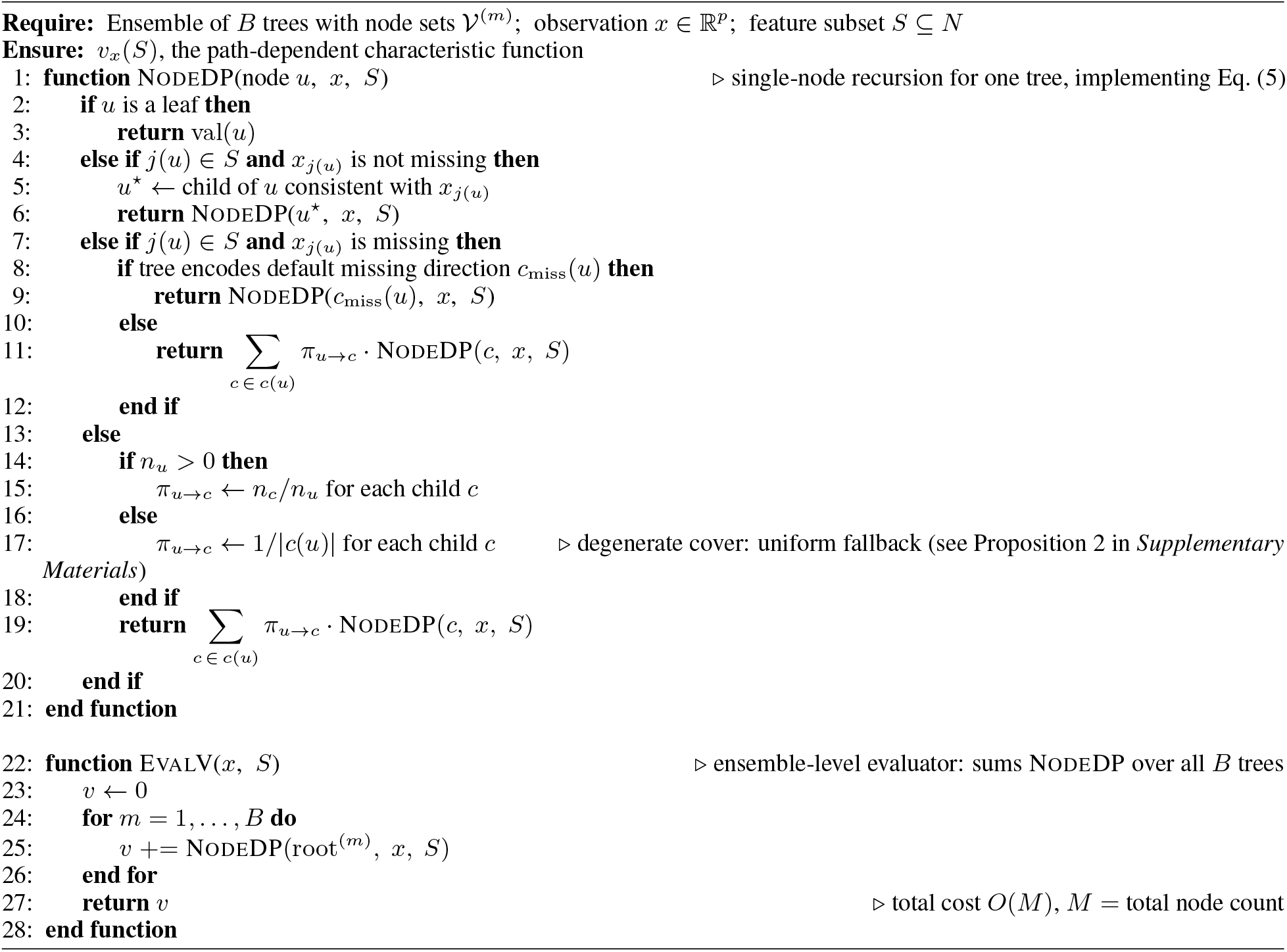

### 2.4 TreeOwen: Monte Carlo Approximation for Large Groups

When within-group sizes |*G*_*k*_| are large, exact enumeration of all 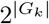 subsets in Step 2 becomes computationally prohibitive. Since Step 1 already controls the number of outer group contexts, the remaining bottleneck lies in the within-group enumeration. The Monte Carlo approximation therefore targets only this inner step, replacing the exact within-group Shapley computation with a permutation-based estimator while keeping the outer aggregation exact. This reduces the inner cost from exponential to linear in |*G*_*k*_| per sample. The procedure is summarized in Algorithm 3.

For a fixed context *C* and group *G*_*k*_, the within-group Shapley value in Eq. (6) admits the permutation representation (Shapley, 1953): for *i* ∈ *G*_*k*_,

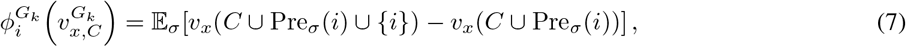

where *σ* is a uniform random permutation of *G*_*k*_ and Pre_*σ*_(*i*) denotes the set of features preceding *i* in *σ*. The expectation is estimated by averaging marginal contributions over sampled permutations, each evaluated via EvalV.

Let *R* denote the number of Monte Carlo units. With antithetic sampling, one unit consists of a pair 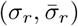. Each *σ*_*r*_ is paired with its reverse 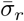, yielding 2*R* permutations in total. This preserves unbiasedness and can reduce variance when the paired contributions are negatively correlated (Proposition 3(ii)). The estimator is

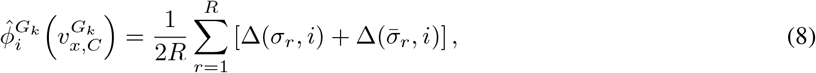

where

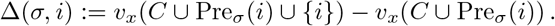

Each permutation requires at most |*G*_*k*_| + 1 evaluations, each costing *O*(*M*), giving per-context cost *O*(*R M* |*G*_*k*_|). For any *R*, the estimator is unbiased; as *R* → ∞, it converges almost surely to the within-group Shapley value

#### Algorithm 2

TreeOwen: Exact Algorithm

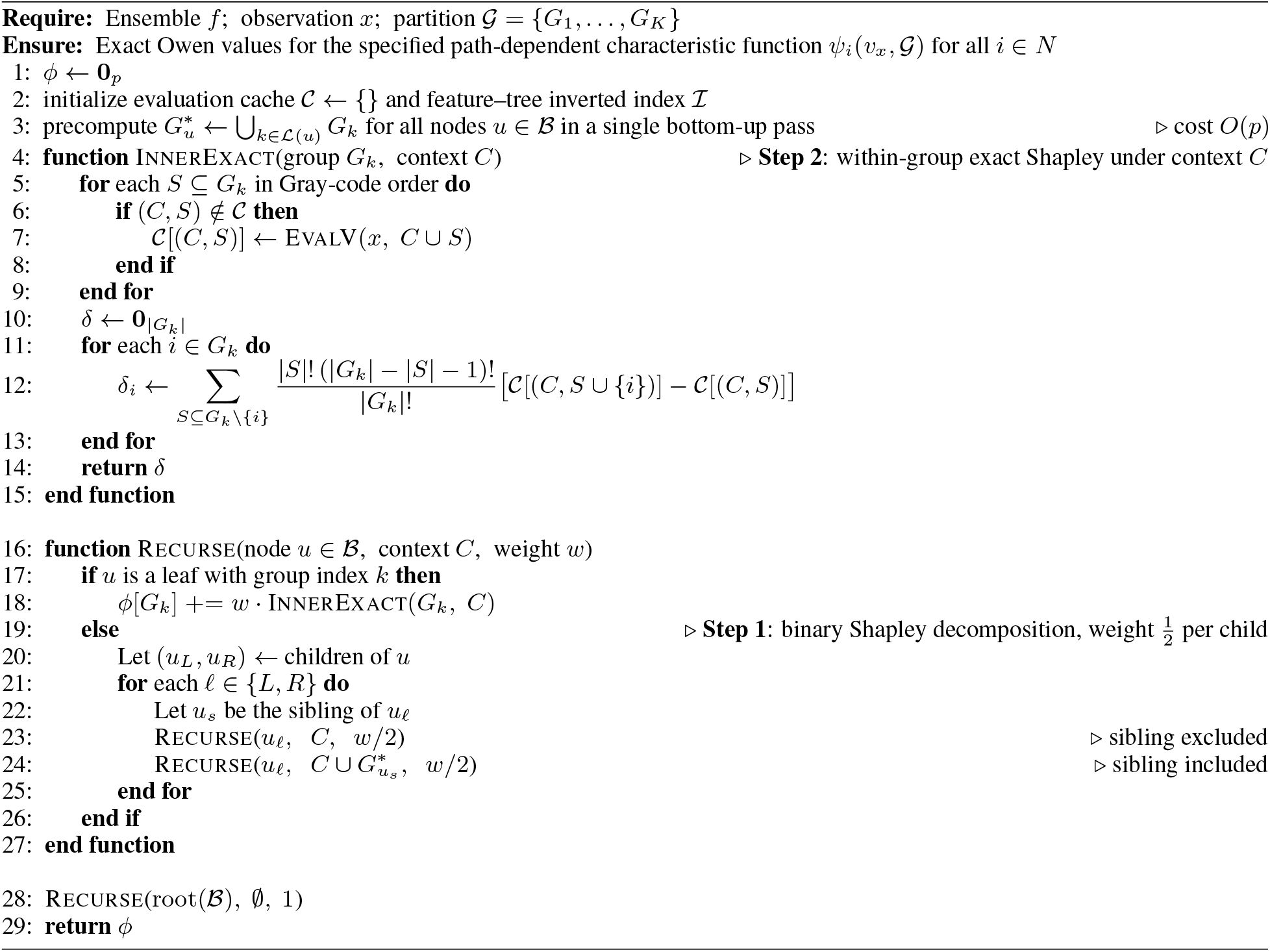

(Proposition 3), with standard error of order *O*(*R*^−1*/*2^). In practice, adaptive stopping is used: sampling continues until the estimated standard error 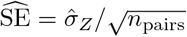 falls below a tolerance *ϵ*, subject to bounds *R*_min_ and *R*_max_. Defaults (*R*_min_ = 32, *R*_max_ = 1024) were chosen as conservative bounds.

Let *d*_*k*_ denote the depth of *G*_*k*_ in ℬ. Since each group is visited under at most 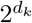 contexts (Lemma 1), the worst-case runtime is

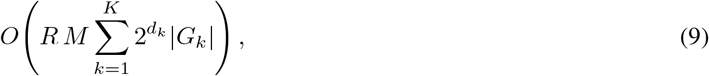

which reduces to

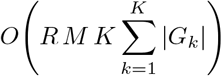

for a balanced tree.

#### Proposition 3

(Statistical Properties of the Antithetic MC Estimator).

*Fix a group G*_*k*_, *an outer context C* ⊆ *N* \ *G*_*k*_, *and define* 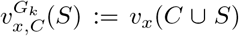 *for S* ⊆ *G*_*k*_. *For any integer R* ≥ 1, *let σ*_1_, …, *σ*_*R*_ *be i*.*i*.*d. uniform random permutations of G*_*k*_ *with reverses* 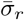, *and let* Δ(*σ, i*) *be as in Eq. (8)*. *The following properties hold for the antithetic estimator* 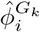 *in Eq. (8) when* `anti = true`; *when* `anti = false`, *unbiasedness and almost sure convergence follow from standard arguments (Hammersley and Handscomb, 1964; Castro et al., 2009; Maleki et al., 2013)*.

i. *(Unbiasedness) For every i* ∈ *G*_*k*_,

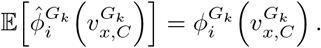
ii. *(Variance formula) Let* Var_*i*_(Δ) := Var_*σ*_[Δ(*σ, i*)]. *Then*

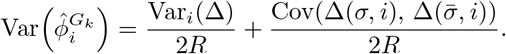

*In particular*, 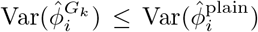 *whenever* 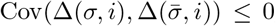, *where* 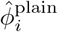 *is the plain MC estimator using* 2*R i*.*i*.*d. draws*.
iii. *(Almost sure convergence)* 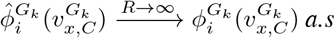.

#### See Proof of Proposition 3 in Supplementary Materials

Proposition 3 concerns the estimator with a prespecified number (*R*) of antithetic pairs. The adaptive stopping rule used in Algorithm 3 is not covered by this proposition, and finite-sample unbiasedness is therefore not claimed for the adaptively stopped estimator. Proposition 3(ii) gives only a conditional variance comparison: reverse-permutation pairing reduces variance relative to (2*R*) independent permutations when the paired marginal contributions have nonpositive covariance. We therefore make no general variance-reduction claim. Recent studies provide further structural explanations for the effectiveness of paired sampling in Shapley-value estimation (Covert and Lee, 2021; Mitchell et al., 2022; Mayer and Wüthrich, 2025; Fumagalli et al., 2026).

### 2.5 Visualization and Importance Measures

TreeOwen provides local visualizations and global importance measures at both the feature and group levels.

#### Local Explanation

*Beeswarm Visualization*. For a dataset of *n* observations, Owen values are computed for each feature and summarized using beeswarm plots analogous to SHAP (Lundberg et al., 2020).

For each feature *i* ∈ *N*, the distribution of Owen values 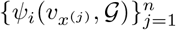 is visualized across observations. Each point corresponds to an observation *j*, positioned horizontally at 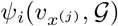 and jittered vertically; its color encodes the feature value 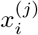. Features are ordered by decreasing mean absolute Owen value,

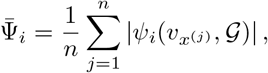

so that the most influential features appear at the top.

At the group level, the Owen value of group *G*_*k*_ for observation *j* is

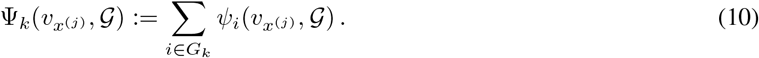

By the efficiency property of the Owen value (Owen, 1977),

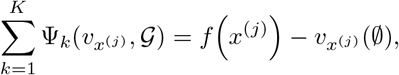

so group-level attributions exactly partition the prediction relative to the path-dependent baseline. The group-level beeswarm plot displays the distribution of 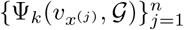, with points colored by a group-level summary statistic. By default, 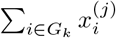 is used, though alternatives (e.g., the mean 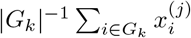 or a user-defined function) may also be specified. Groups are ordered by decreasing mean absolute group attribution,

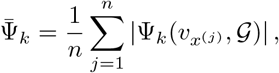

placing the most influential groups at the top.

#### Global Importance Measures

Beyond local attributions, global importance is obtained by aggregating attributions across observations, in the spirit of additive global importance measures such as SAGE (Covert et al., 2020). Unlike SAGE, which defines global importance through a predictive-value game, the proposed measures aggregate local Owen attributions obtained from a prespecified group-structured game. The global feature importance of feature *i* ∈ *N* is defined as

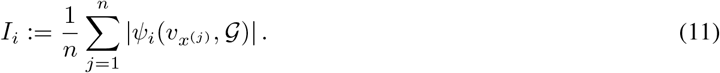

##### Algorithm 3

TreeOwen: Monte Carlo Algorithm

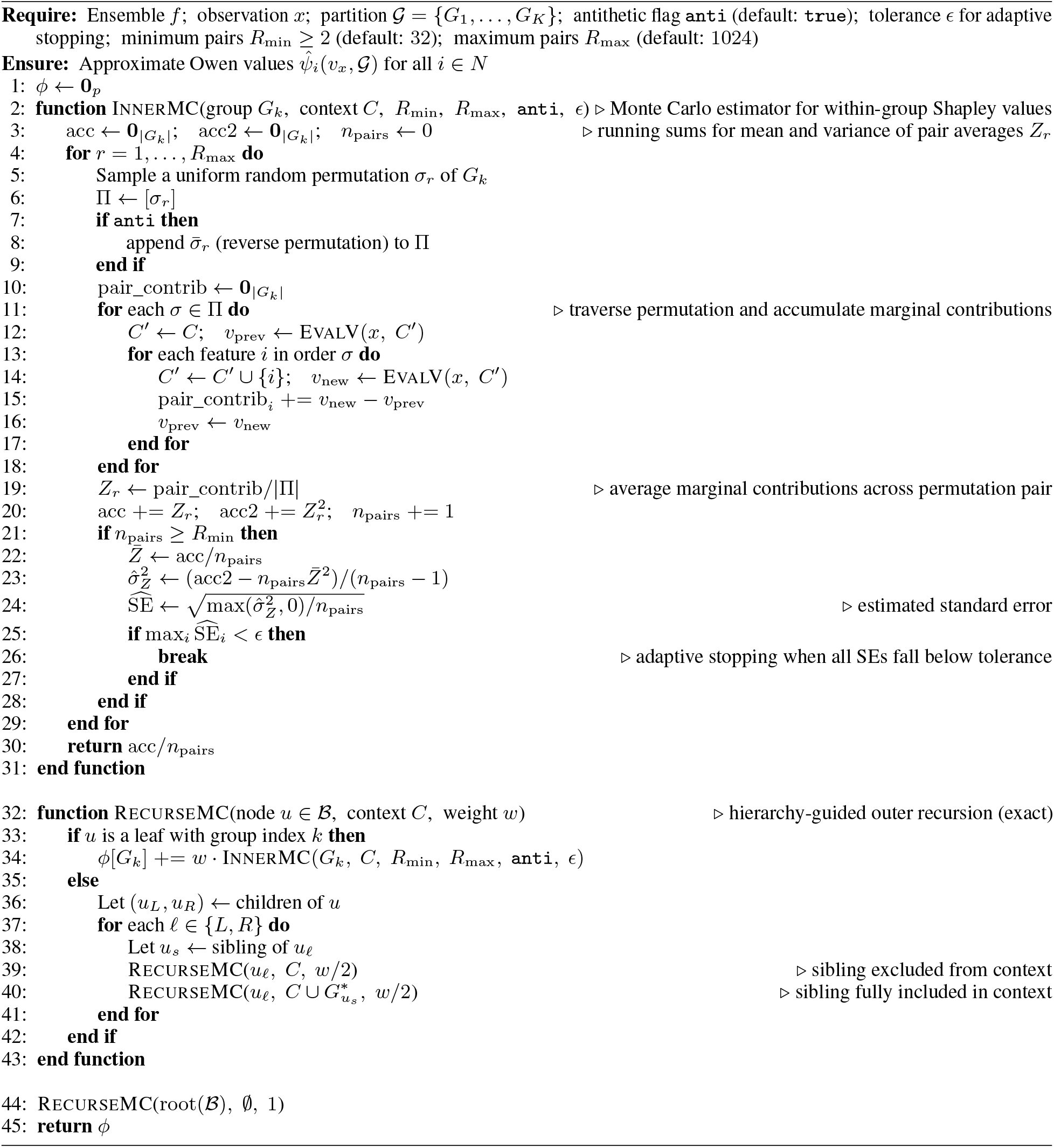

The global group importance of group *G*_*k*_ is

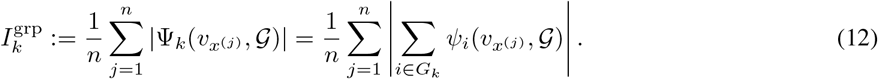

Note that 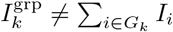 in general, since absolute values do not distribute over sums when within-group contributions have mixed signs. Thus, 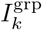 captures the net group contribution rather than the sum of individual magnitudes.

### 2.6 Software: TreeOwen

The algorithms in Algorithms 1–3 are implemented in TreeOwen, an open-source R package available at https://github.com/hk1785/treeowen, compatible with XGBoost (Chen and Guestrin, 2016), LightGBM (Ke et al., 2017), and Ranger (Wright and Ziegler, 2017). Core routines are implemented in C++ with optional multi-core parallelization and a chunked evaluation strategy for memory efficiency. TreeOwen supports explicit selection of either the exact or Monte Carlo algorithm, as well as an automatic mode based on a user-specified within-group size threshold. Under the default threshold of 15, groups with |*G*_*k*_| ≤ 15 are evaluated using the exact algorithm, whereas groups with |*G*_*k*_| > 15 are evaluated using the Monte Carlo algorithm. The package also provides global importance measures (Eqs. (11)–(12)) and visualization tools for feature- and group-level analyses, enabling structured, multi-resolution interpretation of tree-based ensemble predictions.

## 3 Results

### 3.1 Simulation Designs

To evaluate TreeOwen, we conduct controlled simulations to assess numerical accuracy and computational efficiency against exact enumeration baselines (Section 2.2). Both regression and binary classification settings were considered using gradient boosting machines (Friedman, 2001) (XGBoost (Chen and Guestrin, 2016) and LightGBM (Ke et al., 2017)) and random forests (Breiman, 2001) (Ranger (Wright and Ziegler, 2017)). Gradient boosting machines used learning rate 0.2, maximum depth 3, feature subsampling rate 0.5, *ℓ*_2_ regularization, and 500 iterations, while random forests used 500 trees, 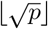 features per split, maximum depth 3, and minimum node size 10 (regression) or 5 (classification). All hyperparameters were fixed across simulation configurations. Table 1 summarizes the complete simulation design, including the data-generating process, group structures, learners, competing methods, Monte Carlo settings, and evaluation metrics.

**Table 1:** Summary of the simulation design.

| Component | Setting |
| --- | --- |
| Data-generating process | A latent-factor model for the $p$ features: $X_j = Z + U_{k(j)} + E_j$ for $j = 1, \dots, p$, where $k(j)$ denotes the group containing feature $j$, and the global factor $Z$, the group factors $U_k$, and the idiosyncratic noises $E_j$ are mutually independent $\mathcal{N}(0, 1)$ variables. Features within a group thus share $U_k$, whereas cross-group dependence arises only through $Z$ |
| Response | Continuous: $Y = \mathbf{X}^\top \beta + \varepsilon$ with independent coefficients $\beta_j \sim \mathcal{N}(0, 1)$ and noise $\varepsilon \sim \mathcal{N}(0, 1)$ (regression); Binary: $\mathbb{P}(Y = 1 \mid \mathbf{X}) = \text{logit}^{-1}(\mathbf{X}^\top \beta)$ (classification) |
| Observations | One dataset of $n = 100$ per configuration, with Owen values computed per observation |
| Group structures | Equal-size ($p = 10K$) and unequal-size ($p = K(K + 1)/2$) designs, with $K = 2, \dots, 20$ groups |
| Learners | XGBoost and LightGBM (learning rate 0.2, depth 3, feature subsampling 0.5, $\ell_2$ regularization, 500 iterations); Ranger (500 trees, $\lfloor \sqrt{p} \rfloor$ features per split, depth 3, minimum node size 10 for regression and 5 for classification) |
| Competing methods | Model-Agnostic Enumeration, Tree-Aware Enumeration, TreeOwen (Exact), and TreeOwen (MC) |
| Monte Carlo settings | 64 inner permutations, bounds $[32, 1024]$, target SE $10^{-3}$ |
| Accuracy | Pearson correlation, agreement within relative tolerance $10^{-3}$, and comparative boxplots |
| Efficiency | Average per-observation runtime; runs beyond five hours reported as 300+ minutes |
| Environment | Single-core C++ on a Mac Studio (M1 Ultra, 64 GB, macOS Tahoe 26.3.1) |

i. *Data-generating process*. The feature vector **X** ∈ ℝ^*p*^ was generated from a latent-factor model inducing within-group correlation and between-group heterogeneity. Each feature *X*_*j*_, *j* = 1, …, *p*, belonging to group *k*(*j*) was drawn as

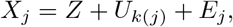

where *Z* ~ *N* (0, 1) is a global factor, *U*_*k*_ ~ *N* (0, 1) is a group-specific factor, and *E*_*j*_ ~ *N* (0, 1) is idiosyncratic noise, all mutually independent. This induces a block-correlation structure: features within a group share *U*_*k*_, while cross-group dependence arises only through *Z*. The response was generated from

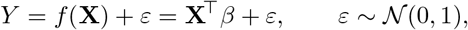

with *β*_*j*_ ~ *N* (0, 1) for regression. For binary classification, *f* (**X**) was passed through the logistic function,

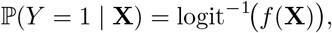

and *Y* was sampled accordingly. For each simulation configuration, one dataset containing (*n* = 100) observations was generated, and Owen values were computed separately for all observations. The reported accuracy measures summarize the resulting observation-level attribution comparisons, while computational efficiency is reported as the average per-observation runtime. The simulations were designed primarily to evaluate attribution accuracy and computational performance at the observation level, rather than predictive-model estimation as a function of sample size.
ii. *Group structures*. Two designs were considered. In the *equal-size* case, *K* groups each contained ten features (*p* = 10*K*), with *K* = 2, …, 20. In the *unequal-size* case, group *k* contained *k* features (*k* = 1, …, *K*), giving *p* = *K*(*K* + 1)*/*2, with *K* = 2, …, 20.
iii. *Competing methods*. Four methods were evaluated: (i) Model-Agnostic Enumeration; (ii) Tree-Aware Enumeration; (iii) TreeOwen (Exact); and (iv) TreeOwen (MC), using *n*_inner_ = 64, target standard error 10^−3^, and bounds [32, 1024].
iv. *Evaluation metrics*. Numerical accuracy was evaluated using Pearson correlation, the percentage of Owen values agreeing within relative tolerance 10^−3^, and comparative boxplots. Computational efficiency was measured by average per-observation runtime using a single-core C++ implementation on a Mac Studio (M1 Ultra, 64 GB, macOS: Tahoe 26.3.1). Configurations exceeding five hours were deemed infeasible and reported as 300+ minutes. Accuracy comparisons were conducted only when both methods produced results.

### 3.2 Simulation Results

We present simulation results on numerical accuracy and computational efficiency for regression and classification tasks using three learners: XGBoost (Chen and Guestrin, 2016), LightGBM (Ke et al., 2017), and Ranger (Wright and Ziegler, 2017), comparing four methods—Model-Agnostic Enumeration, Tree-Aware Enumeration, TreeOwen (Exact), and TreeOwen (MC). The relative performance of these methods is consistent across learners. Thus, for brevity, we present only the results for XGBoost in the main text (Figure 1 for numerical accuracy and Figure 2 for computational efficiency), while results for LightGBM and Ranger are provided in the Supplementary Materials (Figures S1–S2 and S3–S4, respectively).

**Figure 1.**
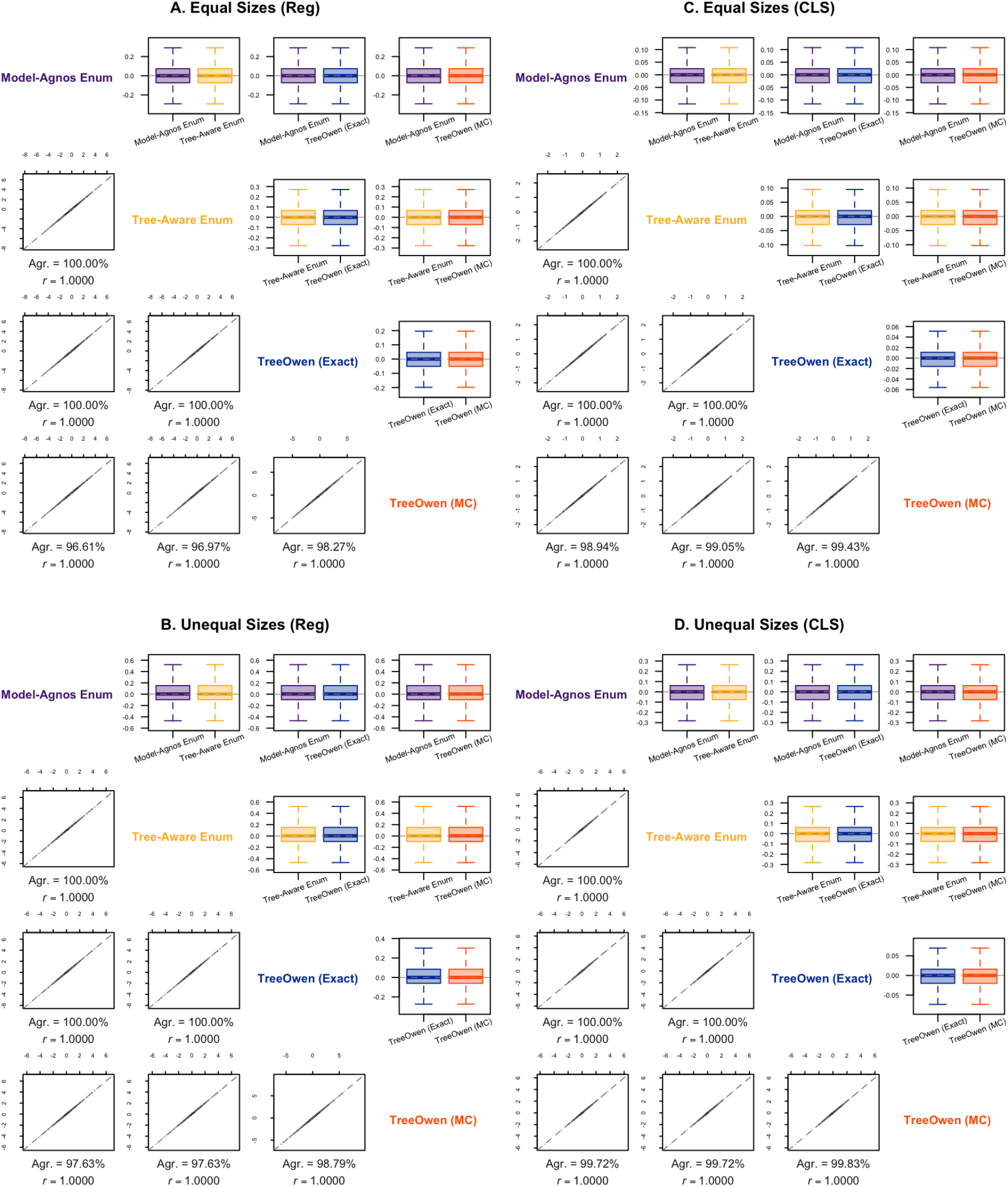
: Results on numerical accuracy, comparing four methods—Model-Agnostic Enumeration (denoted as Model-Agnostic Enum), Tree-Aware Enumeration (denoted as Tree-Aware Enum), TreeOwen (Exact), and TreeOwen (MC)— using three pairwise metrics: (i) Pearson correlation (denoted as *r*), (ii) the percentage of Owen values agreeing within a relative tolerance of 10^−3^ (denoted as Agr.), and (iii) comparative boxplots of Owen value distributions, where the learner is XGBoost. A. Results based on the equal-size design for the regression task; B. Results based on the unequal-size design for the regression task; C. Results based on the equal-size design for the classification task; D. Results based on the unequal-size design for the classification task. TreeOwen (Exact) achieved 100% agreement with enumeration, while TreeOwen (MC) achieved 96.6%–99.8% agreement under a relative tolerance of 10^−3^.

**Figure 2.**
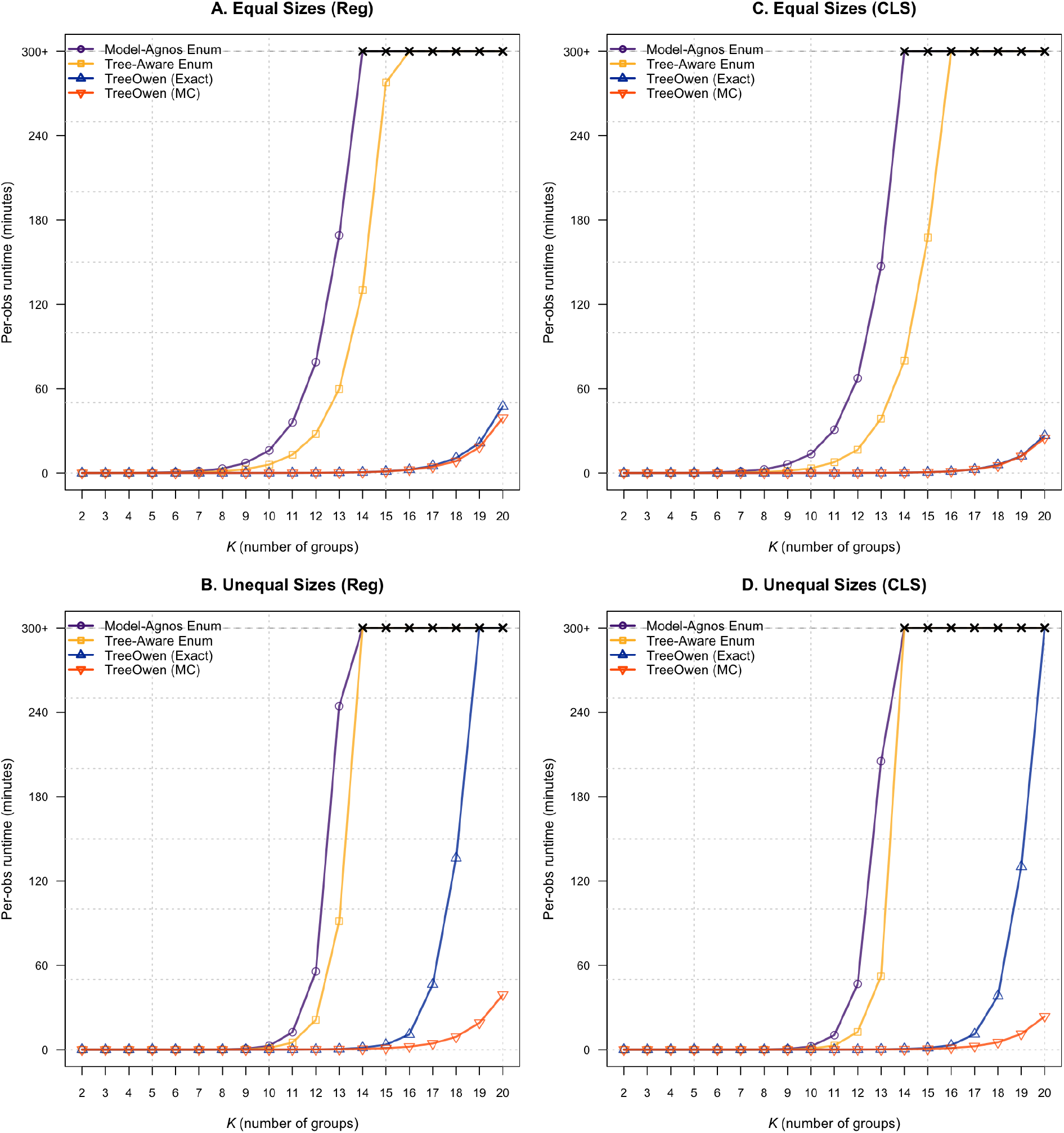
: Results on computational efficiency, comparing four methods—Model-Agnostic Enumeration (denoted as Model-Agnostic Enum), Tree-Aware Enumeration (denoted as Tree-Aware Enum), TreeOwen (Exact), and TreeOwen (MC)—using the average per-observation runtime based on a single-core C++ implementation on a Mac Studio with the M1 Ultra CPU and 64 GB of memory, where the learner is XGBoost. Any configuration whose average per-observation runtime exceeded five hours was considered infeasible and is reported as exceeding 300 minutes (300+). A. Results based on the equal-size design for the regression task; B. Results based on the unequal-size design for the regression task; C. Results based on the equal-size design for the classification task; D. Results based on the unequal-size design for the classification task. Runs exceeding five hours are displayed as 300 + min.

Across all configurations, TreeOwen (Exact) reproduces the reference Owen values from Model-Agnostic and Tree-Aware Enumeration with Pearson correlation of 1.000, 100% agreement under relative tolerance 10^−3^, and identical distributions across both group designs and tasks; see Figures 1, S1, and S3. These results confirm the exactness guaranteed by Theorem 1. TreeOwen (Exact) also achieves substantially lower runtime than both enumeration baselines, consistent with Theorem 2. Both enumeration methods exceeded the five-hour limit at *K* = 14–16 (equal-size) and *K* = 13–14 (unequal-size; max_*k*_ |*G*_*k*_| = 13–14); see Figures 2, S2, and S4. Although Tree-Aware Enumeration is somewhat faster, both methods exhibit exponential runtime growth. In contrast, TreeOwen (Exact) completed all equal-size configurations efficiently and all unequal-size configurations up to *K* = 18–20, with runtime increasing beyond this as within-group enumeration grows exponentially in max_*k*_ |*G*_*k*_|.

TreeOwen (MC) closely matches the reference Owen values from both enumeration methods and TreeOwen (Exact), with perfect or near-perfect Pearson correlation and nearly identical distributions; see Figures 1, S1, and S3. These results demonstrate high empirical accuracy under the evaluated simulation settings. The underlying fixed-budget estimator is unbiased for any prespecified *R*→ ∞ and converges almost surely as *R* by Proposition 3, whereas the adaptive stopping procedure used here is assessed empirically. Agreement under tolerance 10^−3^ ranges from 96% to 99%, reflecting residual Monte Carlo error. TreeOwen (MC) completed all configurations—including those with large groups where TreeOwen (Exact) becomes costly—within a few hours, consistent with the polynomial runtime bound in Eq. (9); see Figures 2, S2, and S4.

These findings motivate the default switching rule in TreeOwen: the exact algorithm is used when |*G*_*k*_| ≤ 15, and the Monte Carlo algorithm otherwise. As noted in Section 2.6, optional multi-core parallelization is also supported.

### 3.3 Real Data Application

This section presents the practical utility of TreeOwen using gut metagenomic data to study the efficacy of cancer immunotherapy in metastatic melanoma patients. The data were compiled in the meta-analysis of Limeta et al. (2020), integrating five independent studies (Matson et al., 2018, Frankel et al., 2017, Gopalakrishnan et al., 2018, Routy et al., 2018, Peters et al., 2019). As a quality control step, subjects with fewer than 2,000 total reads and amplicon sequence variants with mean relative abundance lower than 10^−5^ were excluded. To mitigate study-specific batch effects, the nonparametric correction method ConQuR (Ling et al., 2022) was applied, yielding batch-corrected count data for downstream analysis. The resulting count data were normalized to relative abundances. Taxonomic annotations were standardized, and features were organized into genus-level groups and species-level features. The final dataset comprised 219 patients (145 non-responders and 74 responders) treated with immune checkpoint inhibitors, with 190 microbial species organized into 18 genera used as the coalition structure in TreeOwen.

For predictive modeling, XGBoost (Chen and Guestrin, 2016), LightGBM (Ke et al., 2017), and Ranger (Wright and Ziegler, 2017) were compared using ten-fold cross-validation. The responder base rate was 74*/*219 = 0.338. The cross-validated AUC and AUPRC (mean ± SD across folds) were AUC: 0.863 ± 0.130 and AUPRC: 0.835 ± 0.157 for XGBoost, AUC: 0.878 ± 0.129 and AUPRC: 0.853 ± 0.145 for LightGBM, and AUC: 0.875 ± 0.118 and AUPRC: 0.848 ± 0.130 for Ranger, respectively. LightGBM had the numerically highest mean AUC and AUPRC and was therefore selected for the attribution analysis. The models were trained as follows. LightGBM and XGBoost were configured with a learning rate of 0.05, maximum tree depth of 3, row and feature subsampling rates of 0.8, and up to 1000 boosting iterations. The optimal number of iterations was determined via ten-fold cross-validation with early stopping after 50 rounds. Random forests were trained with 1000 trees, with candidate features per split tuned via out-of-bag error, and a minimum node size of 5. For TreeOwen, the algorithm automatically switched between exact and Monte Carlo computation based on group size: groups with at most 15 features (|*G*_*k*_| ≤ 15) were evaluated exactly, and larger groups (|*G*_*k*_| > 15) via Monte Carlo, matching the package default of Section 2.6. In this dataset, 15 genera were evaluated exactly and 3 via Monte Carlo. The Monte Carlo procedure used 128 inner permutation samples, adaptively controlled between 64 and 1024 to achieve a target standard error of 10^−4^. These settings are stricter than the simulation defaults (*n*_inner_ = 64, bounds [32, 1024], target SE 10^−3^) because the real-data attributions are interpreted individually and therefore warrant tighter standard-error control. Table 2 summarizes the cohort, the coalition structure, the cross-validated predictive performance of the three learners, and the exact and Monte Carlo settings used by TreeOwen.

**Table 2:**
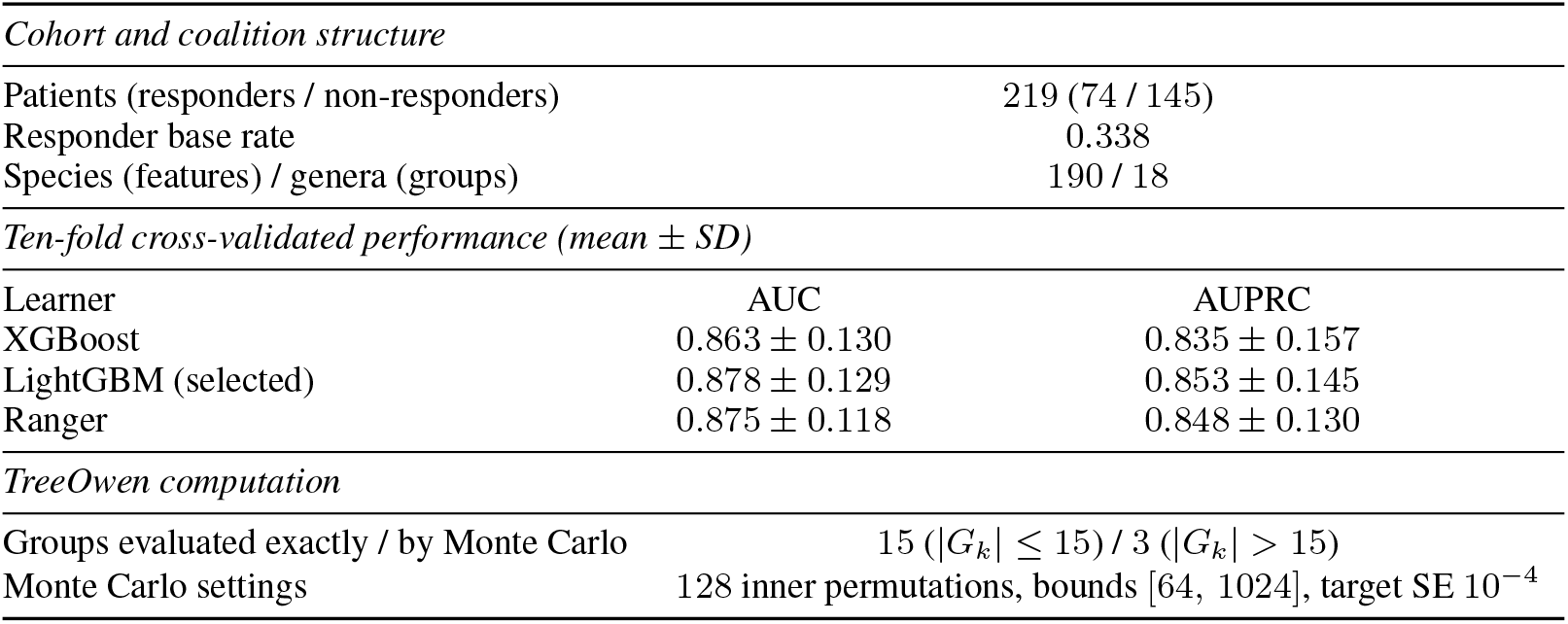
Summary of the real data application. AUPRC is to be read against the responder base rate of 0.338.

Results are presented in Figure 3 using beeswarm plots for the 10 most important genera and their nested species, restricted to the top 100 species by feature importance and ordered by decreasing group importance and then feature importance within each group. Results for all genera and species are provided in the Supplementary Materials (Figures S5–S7). For the three genera evaluated by Monte Carlo, attribution results across 10 independent random seeds were highly stable, with all pairwise Pearson correlations exceeding 0.99. The remaining 15 genera were evaluated exactly and are therefore deterministic.

**Figure 3.**
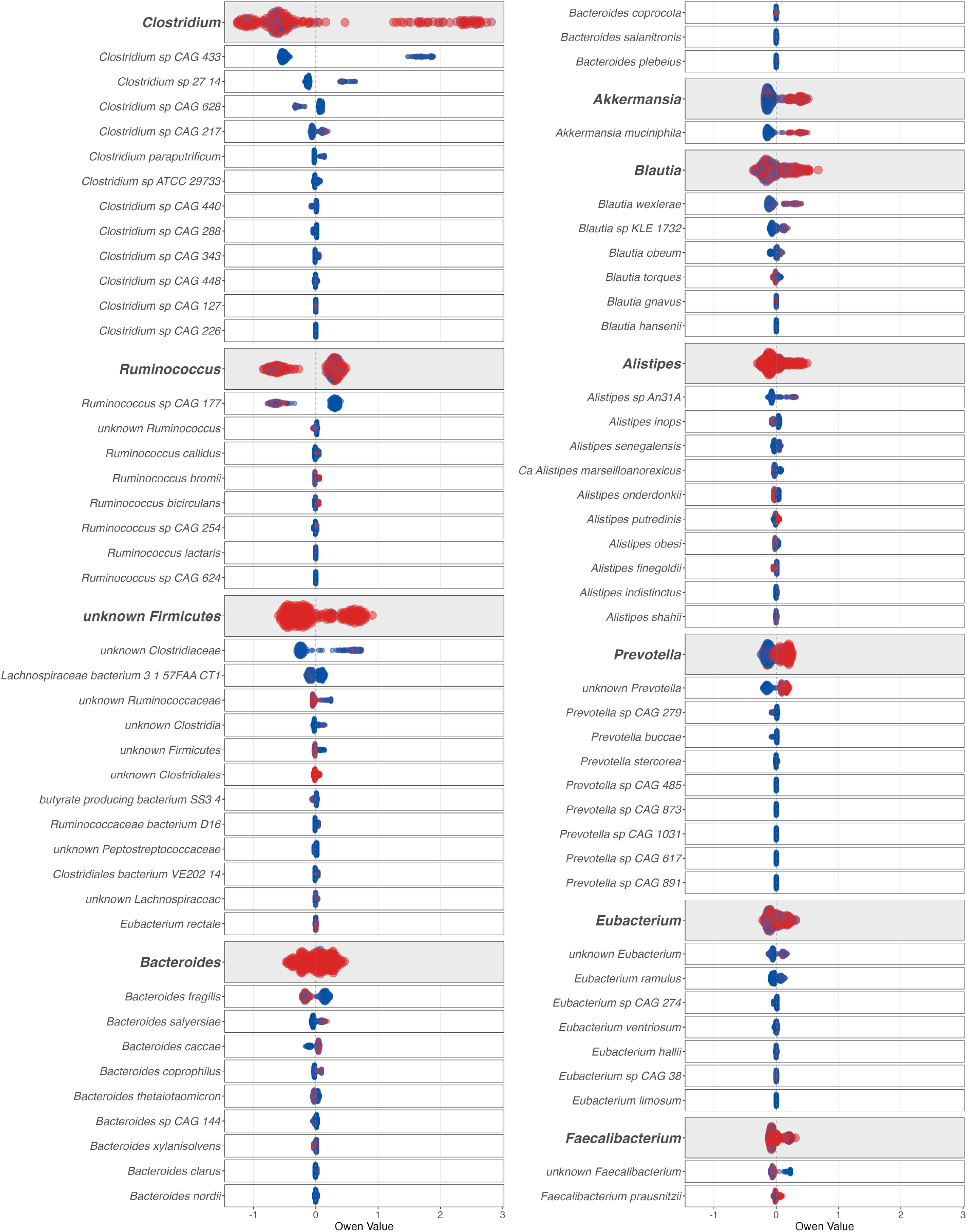
: Results from a real data application using gut metagenomic data to study the efficacy of cancer immunotherapy in metastatic melanoma patients. Beeswarm plots display Owen values for the 10 most important genera (groups, bold italic) and their nested species (features, italic), restricted to the top 100 species by feature importance and ordered by decreasing group importance and then feature importance within each group. Each row shows the distribution of Owen values across patients, with point color indicating relative abundance (blue = low, red = high). ^∗^*unknown* indicates taxa whose formal taxonomic names or positions remain unassigned or unresolved. The selected LightGBM model achieved a cross-validated AUC of 0.878 and AUPRC of 0.853.

We focus on taxa exhibiting high importance and consistent directional effects across patients, while avoiding over-interpretation of *unknown* taxa. Taxa associated with reduced response include *Ruminococcus sp. CAG 177, Bacteroides fragilis*, and *Bacteroides thetaiotaomicron*, consistent with prior studies linking certain *Bacteroides* species to reduced immunotherapy response. In contrast, taxa associated with improved response include the genera *Prevotella, Akkermansia*, and *Blautia*, and species such as *Akkermansia muciniphila, Faecalibacterium prausnitzii*, and *Blautia wexlerae*, consistent with evidence on beneficial roles of short-chain fatty acid–producing bacteria.

Overall, TreeOwen recovers biologically meaningful signals consistent with prior studies while revealing finer-grained patterns that are difficult to detect with standard attribution methods.

## 4 Discussion

In this paper, we introduced TreeOwen, a computational framework comprising exact and Monte Carlo algorithms for efficiently computing Owen values in tree-based ensemble models, including gradient boosting machines (Friedman, 2001) (XGBoost (Chen and Guestrin, 2016) and LightGBM (Ke et al., 2017)) and random forests (Breiman, 2001) (Ranger (Wright and Ziegler, 2017)). The exact algorithm decomposes the computation into hierarchy-guided aggregation across groups and tree-aware dynamic programming within each outer group context, avoiding explicit outer subset enumeration and substantially reducing the combinatorial burden while preserving exactness. The Monte Carlo algorithm further improves scalability for large within-group sizes by replacing exhaustive inner enumeration with a permutation-based estimator that is unbiased for any prespecified sampling budget and converges almost surely as the sampling budget increases.

Through simulation experiments, the exact algorithm achieved perfect numerical accuracy (perfect agreement and Pearson correlation), while the Monte Carlo algorithm closely approximated reference values. In terms of computational efficiency, the exact algorithm substantially outperforms enumeration baselines but becomes costly for large within-group sizes; the Monte Carlo algorithm handles all configurations within practical time limits. The default switching rule, using the exact algorithm when the group size is at most 15 and the Monte Carlo algorithm otherwise, is recommended in practice.

TreeOwen further provides global importance measures and visualization tools for structured, multi-resolution interpretation. Its practical utility was demonstrated on gut metagenomic data from a cancer immunotherapy study, where results were broadly consistent with established biological findings while revealing finer-grained patterns difficult to detect with standard attribution methods.

TreeOwen is intended for settings in which a scientifically meaningful grouping is specified *a priori*, particularly when interactions extend across group boundaries. More generally, the grouping is treated as external domain information that defines the explanation question rather than the prediction rule; TreeOwen neither infers the grouping from the trained model nor assumes that it is represented internally by the trained model. The choice of characteristic function provides a further direction for extending the framework. TreeOwen adopts the path-dependent construction of TreeSHAP (Lundberg et al., 2020), which incorporates feature-dependence information encoded in the split structure and node cover statistics of the trained tree ensemble, without requiring explicit estimation of the conditional feature distribution. This choice enables deterministic, sampling-free computation that is exact for the path-dependent characteristic function while retaining the computational efficiency of tree-specific evaluation. Although the resulting target does not, in general, coincide with the true conditional expectation, it provides a practical balance among dependence-aware attribution, exact computation for the specified game, and computational tractability. Accordingly, any faithfulness claim is limited to the specified path-dependent game: TreeOwen exactly allocates the trained-model prediction relative to the path-dependent baseline, but does not claim to recover the generally unknown observational conditional expectation or causal effects. Marginal or interventional formulations, as used in KernelSHAP and interventional TreeSHAP, average missing features over a background distribution without conditioning them on the observed features, thereby not preserving the dependence between observed and missing features in the coalition evaluation and defining a different attribution target (Lundberg and Lee, 2017; Lundberg et al., 2020; Sundararajan and Najmi, 2020; Janzing et al., 2020; Chen et al., 2020). Explicitly conditional formulations seek to preserve this dependence through an estimated conditional distribution, but require additional distributional modeling and computation (Aas et al., 2021). These characteristic functions offer complementary attribution targets, and extending TreeOwen to support and compare them systematically represents an important direction for future research.

Recent studies have introduced several useful ideas for improving the computation of Shapley values and Shapley interactions for tree-based models. Linear TreeSHAP (Yu et al., 2022) uses node aggregation and telescoping updates to avoid some repeated path-wise computations while retaining the exact path-dependent attribution target. Zern et al. (2023) consider exact interventional Shapley value and interaction computation for piecewise-linear regression trees, thereby extending the class of tree models that can be accommodated. Quadrature-TreeSHAP (Wettenstein et al., 2026) develops a numerical-quadrature formulation for path-dependent Shapley values and arbitrary-order interactions that may offer computational and numerical advantages, particularly for deep trees or higher-order interactions. Nadel and Wettenstein (2026) recast decision-tree computations through Boolean-logic representations and propose a unified framework for exact path-dependent and background SHAP values and interactions. Collectively, these studies suggest several complementary directions for standard feature-level Shapley attribution and interaction analysis. TreeOwen instead focuses on the structured Owen allocation induced by an *a priori* partition, providing coherent attribution both across groups and among the features within each group. Whether and to what extent the computational ideas underlying these recent methods can be adapted effectively to this two-resolution allocation remains an open question. Nevertheless, investigating such adaptations may be worthwhile and may reveal useful opportunities to improve the efficiency and scope of Owen-value computation in future work.

Recent work on paired and structured sampling may also inform the Monte Carlo component of TreeOwen. Mitchell et al. (2022) examine antithetic, orthogonal, and quasi-Monte Carlo designs for permutation-based Shapley estimation, illustrating that the choice and arrangement of sampled permutations can affect estimation accuracy. Covert and Lee (2021) study complement pairing in regression-based KernelSHAP, while Mayer and Wüthrich (2025) provide related theoretical analysis of paired-sampling approximations. Fumagalli et al. (2026) further describe a structural interpretation of complement pairing in regression-based Shapley estimation through the odd and even components of the underlying set function. TreeOwen uses a related antithetic construction by pairing each sampled within-group permutation with its reverse. For each feature, the two orderings generate complementary predecessor coalitions within the corresponding within-group game while preserving estimator unbiasedness. The present analysis establishes unbiasedness for any prespecified number of antithetic pairs and almost sure convergence as the number of pairs increases. It does not claim a general variance-reduction guarantee, although variance may be reduced when the paired marginal contributions are negatively correlated. Future work may compare reverse-permutation pairing with alternative structured-sampling designs and investigate whether related ideas can be adapted usefully to the within-group games arising in Owen-value computation.

In conclusion, TreeOwen offers a practical and principled framework for structured, multi-resolution interpretation in settings where features are naturally organized into *a priori* groups, supporting hierarchical attribution at both the feature and group levels. When combined with tree-based ensemble models that capture complex nonlinear relationships and interactions, TreeOwen serves as a robust and flexible tool for diverse scientific applications. Extensions to alternative characteristic functions and more efficient computational and sampling procedures may further broaden its scope, establishing TreeOwen as a practical foundation for group-aware explanation in scientific applications.

## Acknowledgements

The author is grateful to the anonymous reviewers for their thorough reviews and insightful comments, which helped improve the quality of this manuscript.

## Funding

This work was supported by the National Research Foundation of Korea (NRF) grant funded by the Korean government (MSIT) (No. RS-2026-25477838).

## Conflicts of Interest

The author declares no conflicts of interest.

## Data Availability

The software and tutorials are freely available as an R package, treeowen, at https://github.com/hk1785/treeowen. All processed real datasets used in this study are included in the treeowen package as example data.

## Supplementary Materials

Additional supporting information, including the proofs of Propositions 1–3, Lemma 1, and Theorems 1 and 2, as well as Figures S1–S7, is available online in the Supplementary Materials section.

## Notes

### Competing Interest Statement

The authors have declared no competing interest.

