## Supplementary Materials for "Efficient Game-Theoretic Explanations for Tree-Based Ensembles via Owen Values"

**Hyunwook Koh**

****

**Proposition 1** (Owen versus summed Shapley)**.** *The summed Shapley attribution* $\psi_i^{\mathrm{S}} = \phi_i$ *satisfies (i) group-level efficiency and (iv) additivity, but does not in general satisfy (ii) feature-level efficiency, (iii) symmetry, or (v) the null-player property.*

**Proof of Proposition 1.** Write $\psi_i^{\mathrm{S}}(v_x) = \phi_i(v_x)$ for the summed Shapley attribution, with group total $\sum_{i \in G_k} \phi_i(v_x)$, and recall the group-level (quotient) game $\widetilde{v}_x(T) = v_x\big(\bigcup_{k \in T} G_k\big)$.

*Axioms (i) and (iv).* Both are inherited from the Shapley value: $\sum_{i \in N} \phi_i(v_x) = v_x(N) - v_x(\emptyset)$ by efficiency, which gives group-level efficiency (i); and $\phi_i(v_x + w_x) = \phi_i(v_x) + \phi_i(w_x)$ by additivity, which gives (iv).

*Axioms (ii), (iii), and (v) fail in general.* The hypotheses of these axioms are stated relative to the group structure: (ii) refers to the quotient game $\widetilde{v}_x$, while (iii) and (v) quantify only over group-complete outer contexts $v_x^U$ with $U \subseteq \mathcal{K} \setminus \{k\}$, in which each other group is present in full or absent in full. The Shapley value, however, also responds to *partial* coalitions of other groups' features, so it need not respect these group-relativized conditions. Two minimal counterexamples suffice.

*(iii) and (v).* Let $N = \{1,2,3,4\}$, $G_1 = \{1,2\}$, $G_2 = \{3,4\}$, and $v_x(\{1,3\}) = 1$ with $v_x(T) = 0$ for all other $T$. Evaluating the Shapley value, the only nonzero marginal contribution for player 1 occurs when 1 is added to $\{3\}$, and similarly (by the symmetry of $v_x$ under the exchange $1 \leftrightarrow 3$, $2 \leftrightarrow 4$) for the other players, giving

$$\phi_1 = \tfrac{1}{12}, \qquad \phi_2 = -\tfrac{1}{12}, \qquad \phi_3 = \tfrac{1}{12}, \qquad \phi_4 = -\tfrac{1}{12}.$$

Player 1 is a null player *relative to the group structure*: for every $S \subseteq G_1 \setminus \{1\}$ and every $U \subseteq \{2\}$ (context $\emptyset$ or $G_2$) one has $v_x^U(S \cup \{1\}) = v_x^U(S) = 0$, because the coalition $\{1,3\}$ requires the *partial* presence of $G_2$ (namely 3 without 4), which no group-complete context realizes. Yet $\phi_1 = \frac{1}{12} \neq 0$, violating the null-player property (v). Likewise, players 1 and 2 are interchangeable across all group-complete contexts (each yields value 0), yet $\phi_1 = \frac{1}{12} \neq -\frac{1}{12} = \phi_2$, violating symmetry (iii). The Owen value gives $\psi_1 = \psi_2 = 0$ in this example.

*(ii).* Let $N = \{1,2,3\}$, $G_1 = \{1,2\}$, $G_2 = \{3\}$, and $v_x(\{1\}) = 1$ with $v_x(T) = 0$ otherwise. Then $\phi_1 = \frac{1}{3}$ and $\phi_2 = -\frac{1}{6}$, so $\sum_{i \in G_1} \phi_i = \frac{1}{6}$. The quotient game satisfies $\widetilde{v}_x(\{1\}) = v_x(G_1) = 0$ and $\widetilde{v}_x(\{1,2\}) = v_x(N) = 0$, so the Shapley value of group 1 in $\widetilde{v}_x$ equals $0 \neq \frac{1}{6}$, violating feature-level efficiency (ii). By construction the Owen value satisfies (ii) with equality. $\square$

**Proposition 2** (Correctness of NodeDP)**.** *Let $T^{(m)}$ be a fitted decision tree with node set $\mathcal{V}^{(m)}$, and let $X$ be drawn from the empirical training distribution. For any feature subset $S \subseteq N$ and observation $x$, the value $V_{\mathrm{root}}(S)$ returned by NodeDP (Algorithm 1 in the main text) equals the path-dependent characteristic function value—the cover-weighted tree traversal of Lundberg et al. (2020), which approximates $\mathbb{E}[f^{(m)}(X) \mid X_S = x_S]$ but does not in general equal it—under the following two conventions:*

*(i)* (Degenerate cover) *If a node $u$ has training-sample cover $n_u = 0$, branch probabilities are set to $\pi_{u \to c} = 1/|c(u)|$ uniformly.*

*(ii)* (Missing observed value) *If $j(u) \in S$ but $x_{j(u)}$ is missing, NodeDP follows the model's default missing direction $c_{\mathrm{miss}}(u)$ if available, and otherwise falls back to cover-weighted averaging (treating $j(u)$ as unknown).*

*In both cases, NodeDP correctly implements the path-dependent characteristic function $v_x(S)$ used throughout the main text, and all correctness results of TreeOwen hold with respect to this $v_x$ under either convention. The allocation results that treat $v_x$ as a supplied characteristic function remain applicable to alternative characteristic functions, provided that EvalV is replaced by a corresponding evaluator.*

**Proof of Proposition 2.**

**Proof.** (i) A node $u$ with $n_u = 0$ was never reached by any training sample, so the formula $\pi_{u \to c} = n_c/n_u$ is undefined (0/0). Setting $\pi_{u \to c} = 1/|c(u)|$ uniformly defines a valid probability distribution over children; NodeDP then returns the unweighted average of the child-subtree values, providing a well-defined fallback for the path-dependent recursion under this convention.

(ii) When $j(u) \in S$ but $x_{j(u)}$ is missing, the deterministic branch cannot be identified. Falling back to cover-weighted averaging treats $j(u)$ as unknown at node $u$, computing $V_u(S) = \sum_{c \in c(u)} \pi_{u \to c} V_c(S)$ using empirical branch proportions, which is consistent with the path-dependent recursion implemented in Algorithm 1 of the main text. When the fitted tree encodes an explicit default missing direction $c_{\mathrm{miss}}(u)$ (supported by XGBoost and LightGBM), following it deterministically propagates the model's own imputation convention and preserves $v_x(N) = f(x)$ when $x$ contains missing values.

In both cases, NodeDP defines a well-formed probability-weighted averaging scheme over leaf values, and all downstream correctness results follow because they depend solely on EvalV returning $v_x(S)$ for arbitrary $S \subseteq N$. □

**Lemma 1** (Context Count and Weight Partition)**.** *Let $\mathcal{B}$ be a balanced binary tree with $K \geq 1$ leaves, and let $d_k$ denote the depth of the leaf corresponding to group $G_k$. For any leaf $G_k$:*

*(i)* (Count) *RECURSE visits $G_k$ under exactly $2^{d_k}$ distinct outer contexts. Hence $2^{d_k} \leq 2^{\lceil \log_2 K \rceil} \leq 2K - 1$.*

*(ii)* (Weight partition) *If $w_C^{(k)}$ denotes the weight assigned to context $C$ when $G_k$ is visited, then $\sum_C w_C^{(k)} = 1$.*

**Proof of Lemma 1.**

**Proof.** (i) When $K = 1$, the single leaf is also the root, so $d_k = 0$ and RECURSE is called once with the empty context; the bound $2^0 = 1 \leq 2(1) - 1 = 1$ holds. For $K \geq 2$, the path from the root to leaf $G_k$ passes through exactly $d_k \geq 1$ internal ancestors. At each ancestor, RECURSE makes two recursive calls to the branch containing $G_k$: once with the sibling subtree excluded from the context and once with it fully included. Since each of the $d_k$ binary inclusion–exclusion decisions is independent, the context upon reaching $G_k$ is determined by a binary string of length $d_k$, yielding exactly $2^{d_k}$ distinct contexts. For a balanced binary tree with $K \geq 2$, $d_k \leq \lceil \log_2 K \rceil$. Let $m = \lceil \log_2 K \rceil$; then $2^{m-1} < K$, so $2^m < 2K$, and since $2^m$ is an integer, $2^m \leq 2K - 1$.

(ii) Proved by induction on the depth of the tree. We show that *for every node $v$ in $\mathcal{B}$, the total weight over all calls* $\textsc{Recurse}(v, \cdot, \cdot)$ *equals* 1; part (ii) is the instance $v = G_k$.

*Base case* (root): $\textsc{Recurse}(\text{root}, \emptyset, 1)$ is called exactly once, giving total weight 1.

*Inductive step*: let $v$ be any non-root node with parent $u_{\text{par}}$, and assume the total weight at $u_{\text{par}}$ equals 1. Each call $\textsc{Recurse}(u_{\text{par}}, C, w)$ issues two calls to $v$, each with weight $w/2$, contributing total weight $w$ to $v$. Summing over all calls to $u_{\text{par}}$ gives total weight at $v$ equal to $\sum w = 1$. □

**Theorem 1** (Correctness)**.** *Let $\mathcal{G} = \{G_1, \ldots, G_K\}$ be a partition of $N$ and let $v_x : 2^N \to \mathbb{R}$ be any characteristic function. For every $i \in G_k$, Algorithm 2 in the main text returns*

$$\sum_C w_C^{(k)} \, \phi_i^{G_k}\big(v_{x,C}^{G_k}\big),$$

*where the contexts $C$ and weights $w_C^{(k)}$ are as specified in Lemma 1, $v_{x,C}^{G_k}(S) := v_x(C \cup S)$ for $S \subseteq G_k$, and $\phi_i^{G_k}$ denotes the Shapley value of player $i$ in the $|G_k|$-player game $(G_k, v_{x,C}^{G_k})$.*

**Proof of Theorem 1.**

**Proof.** Fix a feature $i \in G_k$. By Lemma 1(i), Recurse visits the leaf $G_k$ under exactly $2^{d_k}$ distinct outer contexts. For such a context $C$, let $w_C^{(k)}$ be the weight assigned to $C$ when $G_k$ is visited, that is, the product of the factors $\frac{1}{2}$ accumulated along the root-to-leaf path. Upon reaching $G_k$ under context $C$, Recurse invokes InnerExact$(G_k, C)$ and accumulates $\phi[G_k] \mathrel{+}= w_C^{(k)} \cdot$ InnerExact$(G_k, C)$.

By construction, InnerExact$(G_k, C)$ enumerates all $2^{|G_k|}$ subsets $S \subseteq G_k$, evaluates $v_x(C \cup S)$ exactly via EvalV, and returns, for each $i \in G_k$,

$$\sum_{S \subseteq G_k \setminus \{i\}} \frac{|S|! \, (|G_k| - |S| - 1)!}{|G_k|!} \big[v_x(C \cup S \cup \{i\}) - v_x(C \cup S)\big].$$

With $v_{x,C}^{G_k}(S) := v_x(C \cup S)$, this is exactly the Shapley value $\phi_i^{G_k}(v_{x,C}^{G_k})$ of player $i$ in the game $(G_k, v_{x,C}^{G_k})$ (Shapley, 1953; Eq. (6) in the main text). The value is exact because all subsets are enumerated and each characteristic-function value is computed exactly by EvalV.

Summing the accumulated contributions over the $2^{d_k}$ contexts visited for $G_k$, Algorithm 2 returns

$$\sum_C w_C^{(k)} \, \phi_i^{G_k}\big(v_{x,C}^{G_k}\big),$$

as claimed; by Lemma 1(ii), these weights satisfy $\sum_C w_C^{(k)} = 1$. □

**Theorem 2** (Runtime Complexity)**.** *Under the conditions of Theorem 1, with $\mathcal{B}$ a balanced binary tree and characteristic function values evaluated via* EvalV *(Algorithm 1 in the main text), the total number of* EvalV *calls is at most $(2K-1)\sum_{k=1}^{K} 2^{|G_k|}$, and the total runtime is*

$$O\big(M \cdot K \sum_{k=1}^{K} 2^{|G_k|}\big),$$

*where $M$ is the total node count across all $B$ trees.*

**Proof of Theorem 2.**

**Proof.** By Lemma 1(i), each $G_k$ is visited under at most $2^{d_k} \leq 2K-1$ distinct contexts. For each context, InnerExact evaluates $v_x(C \cup S)$ for all $2^{|G_k|}$ subsets $S \subseteq G_k$; the global cache $\mathcal{C}$ ensures no pair $(C, S)$ is passed to EvalV more than once, so the number of EvalV calls per group per context is at most $2^{|G_k|}$, each at cost $O(M)$. Summing over all groups and contexts yields at most

$$\sum_{k=1}^{K} 2^{d_k} \cdot 2^{|G_k|} \leq (2K-1) \sum_{k=1}^{K} 2^{|G_k|}$$

EvalV calls, and total runtime $O\big(M \cdot K \sum_{k=1}^{K} 2^{|G_k|}\big)$. □

**Proposition 3** (Statistical Properties of the Antithetic MC Estimator)**.** *Fix a group $G_k$, an outer context $C \subseteq N \setminus G_k$, and define $v^{G_k}_{x,C}(S) := v_x(C \cup S)$ for $S \subseteq G_k$. Fix any integer $R \geq 1$. Let $\sigma_1, \ldots, \sigma_R$ be i.i.d. uniform random permutations of $G_k$ with reverses $\bar{\sigma}_r$, and let*

$$\Delta(\sigma, i) := v_x(C \cup \mathrm{Pre}_\sigma(i) \cup \{i\}) - v_x(C \cup \mathrm{Pre}_\sigma(i))\,.$$

*The following properties hold for the antithetic estimator $\hat{\phi}^{G_k}_i$ in Eq. (8) in the main text when* `anti = true`*; when* `anti = false`*, unbiasedness and almost sure convergence follow from standard arguments (Hammersley and Handscomb, 1964; Castro et al., 2009; Maleki et al., 2013).*

*(i)* (Unbiasedness) *For every $i \in G_k$, $\mathbb{E}[\hat{\phi}^{G_k}_i(v^{G_k}_{x,C})] = \phi^{G_k}_i(v^{G_k}_{x,C})$.*

*(ii)* (Variance formula) *Let $\mathrm{Var}_i(\Delta) := \mathrm{Var}_\sigma[\Delta(\sigma, i)]$. Then*

$$\mathrm{Var}\big(\hat{\phi}^{G_k}_i\big) = \frac{\mathrm{Var}_i(\Delta)}{2R} + \frac{\mathrm{Cov}(\Delta(\sigma, i),\, \Delta(\bar{\sigma}, i))}{2R}.$$

*In particular, $\mathrm{Var}(\hat{\phi}^{G_k}_i) \leq \mathrm{Var}(\hat{\phi}^{\mathrm{plain}}_i)$ whenever $\mathrm{Cov}(\Delta(\sigma, i), \Delta(\bar{\sigma}, i)) \leq 0$, where $\hat{\phi}^{\mathrm{plain}}_i$ is the plain MC estimator using $2R$ i.i.d. draws.*

*(iii)* (Almost sure convergence) $\hat{\phi}^{G_k}_i(v^{G_k}_{x,C}) \xrightarrow{R\to\infty} \phi^{G_k}_i(v^{G_k}_{x,C})$ a.s.

**Proof of Proposition 3.**

**Proof.** Throughout, let $Z_r := \frac{1}{2}[\Delta(\sigma_r, i) + \Delta(\bar{\sigma}_r, i)]$ denote the $r$-th antithetic pair average.

*Part (i): Unbiasedness.* By the permutation representation of the Shapley value (Shapley, 1953), for $\sigma$ uniform on $\mathfrak{S}_{G_k}$, $\mathbb{E}[\Delta(\sigma, i)] = \phi^{G_k}_i(v^{G_k}_{x,C})$. Since reversing a uniform permutation yields a uniform permutation, $\bar{\sigma}_r$ is also uniform on $\mathfrak{S}_{G_k}$, so $\mathbb{E}[\Delta(\bar{\sigma}_r, i)] = \phi^{G_k}_i(v^{G_k}_{x,C})$. By linearity of expectation,

$$\mathbb{E}\big[\hat{\phi}^{G_k}_i\big] = \frac{1}{2R}\sum_{r=1}^{R}\Big(\mathbb{E}[\Delta(\sigma_r, i)] + \mathbb{E}[\Delta(\bar{\sigma}_r, i)]\Big) = \phi^{G_k}_i\left(v^{G_k}_{x,C}\right).$$

*Part (ii): Variance formula.* Since $\bar{\sigma}_r$ is a deterministic function of $\sigma_r$, each $Z_r$ is a measurable function of the single independent draw $\sigma_r$, so $Z_1, \ldots, Z_R$ are i.i.d. By bilinearity of covariance and marginal uniformity,

$$\begin{aligned}\mathrm{Var}(Z_r) &= \tfrac{1}{4}\big[\mathrm{Var}_i(\Delta) + \mathrm{Var}_i(\Delta) + 2\,\mathrm{Cov}(\Delta(\sigma_r, i), \Delta(\bar{\sigma}_r, i))\big] \\ &= \tfrac{1}{2}\mathrm{Var}_i(\Delta) + \tfrac{1}{2}\mathrm{Cov}(\Delta(\sigma, i), \Delta(\bar{\sigma}, i)).\end{aligned}$$

Since $\hat{\phi}^{G_k}_i = R^{-1}\sum_{r=1}^{R} Z_r$, we have $\mathrm{Var}(\hat{\phi}^{G_k}_i) = \mathrm{Var}(Z_r)/R$, giving the stated formula. The plain estimator with $2R$ i.i.d. draws has variance $\mathrm{Var}_i(\Delta)/(2R)$, so the difference $\mathrm{Cov}(\Delta(\sigma, i), \Delta(\bar{\sigma}, i))/(2R) \leq 0$ whenever the covariance is non-positive.

*Part (iii): Almost sure convergence.* Since $v_x$ is bounded on the finite set $2^N$, every $\Delta$-value is bounded, so $Z_1, Z_2, \ldots$ are i.i.d. with finite mean $\mathbb{E}[Z_1] = \phi^{G_k}_i(v^{G_k}_{x,C})$. Because $\hat{\phi}^{G_k}_{i,R} = R^{-1}\sum_{r=1}^{R} Z_r$, the strong law of large numbers gives $\hat{\phi}^{G_k}_{i,R} \to \phi^{G_k}_i(v^{G_k}_{x,C})$ almost surely as $R \to \infty$. □

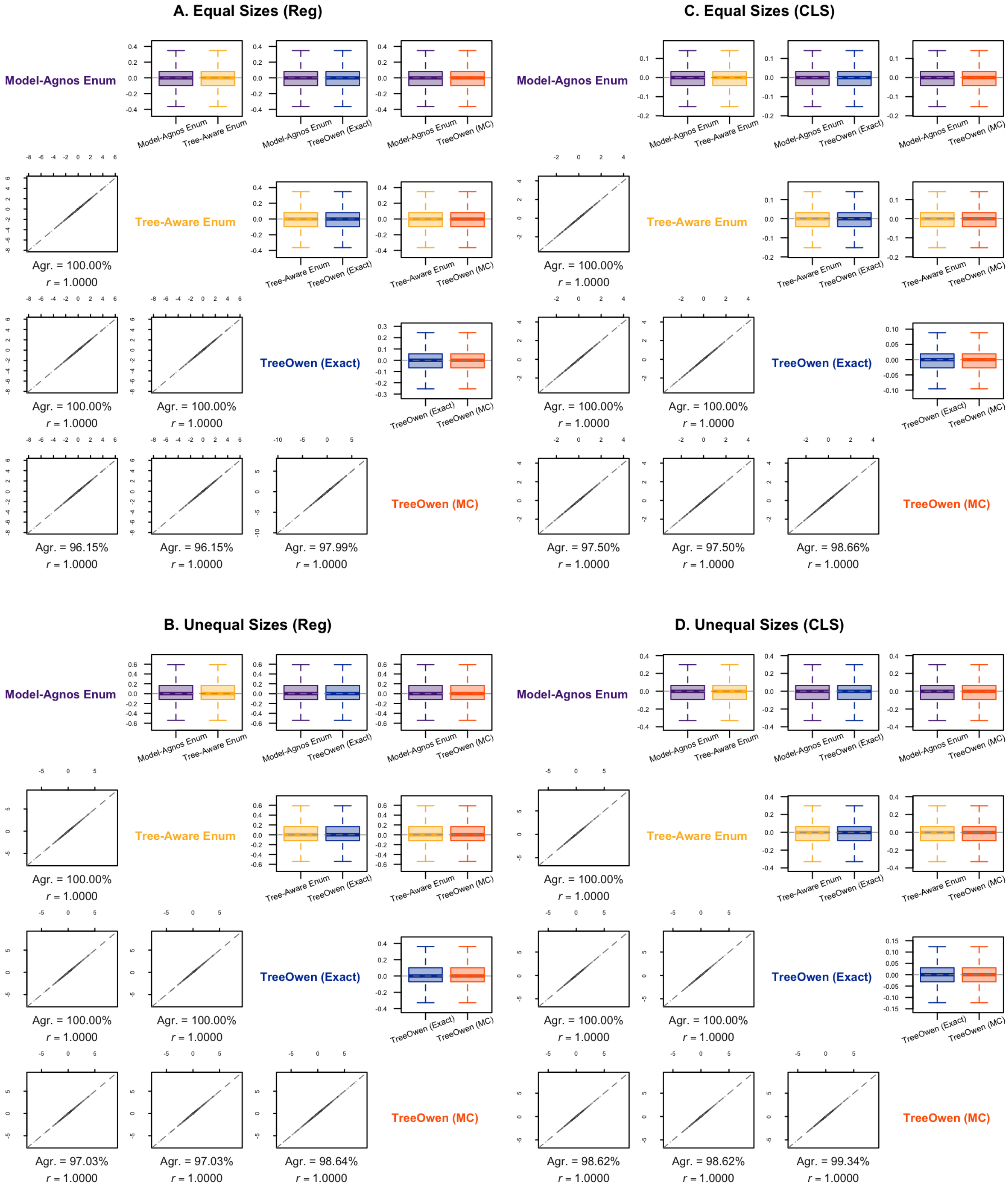


**Fig. S1.** Results on numerical accuracy for LightGBM.

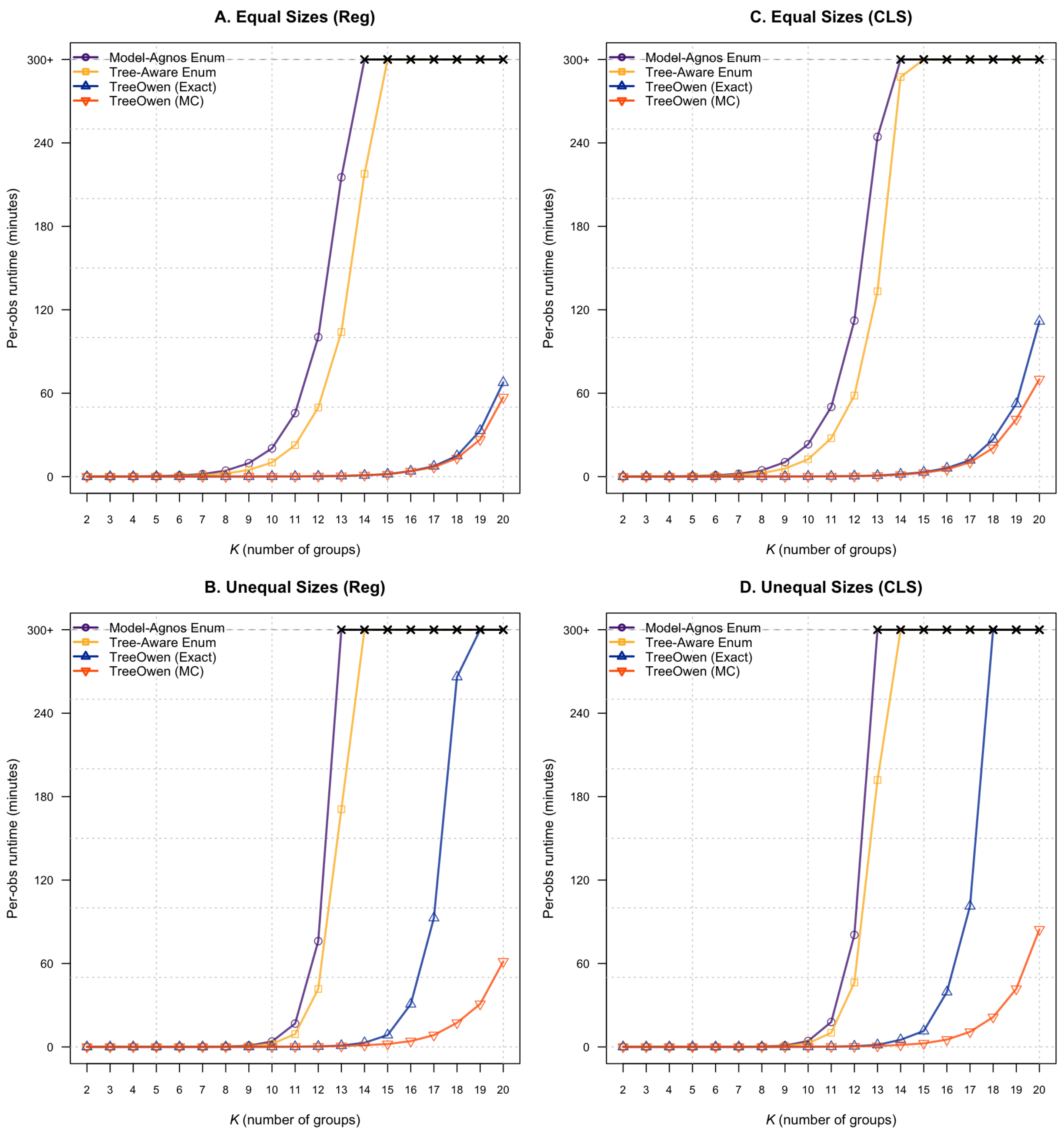


**Fig. S2.** Results on computational efficiency for LightGBM.

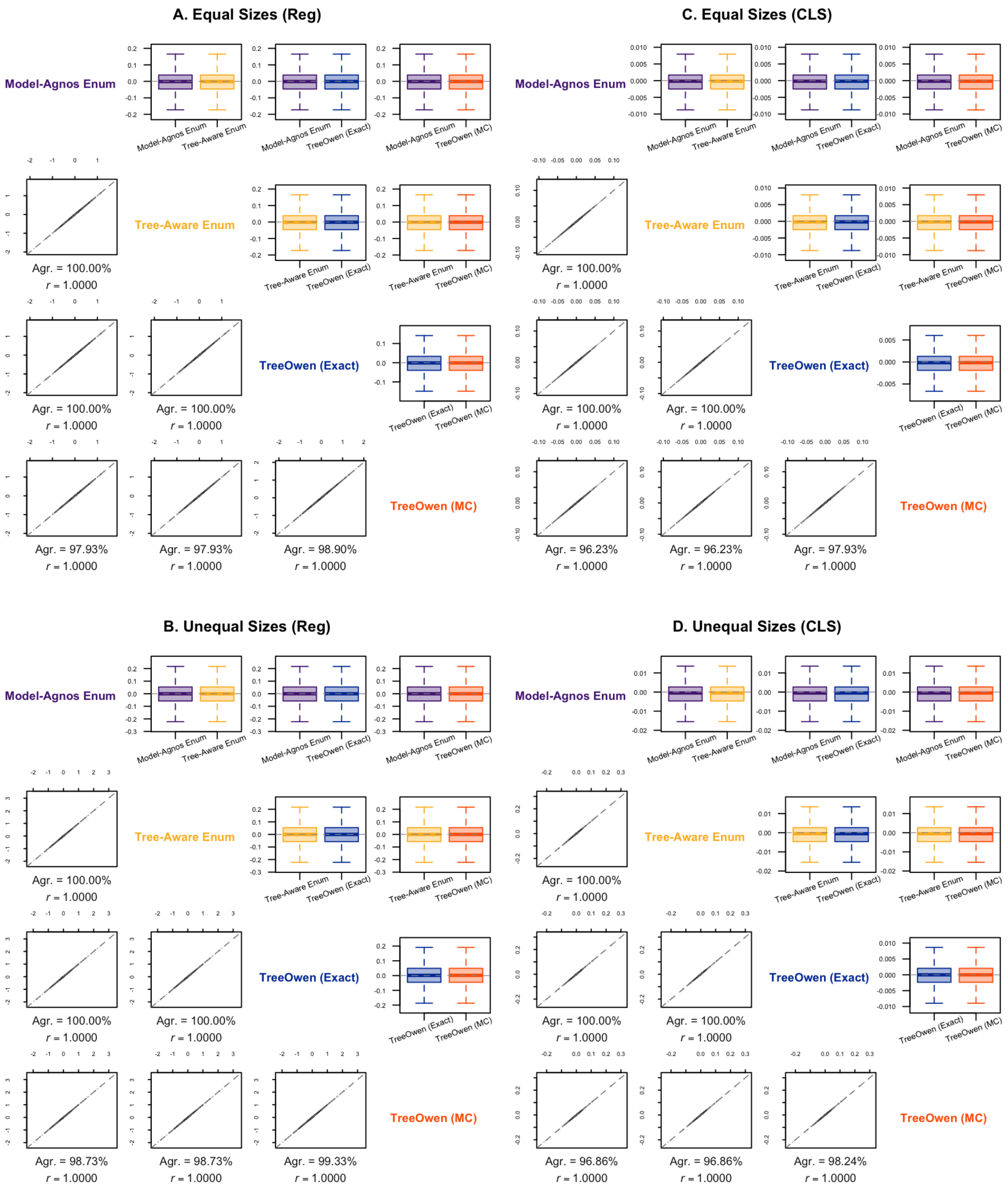


**Fig. S3.** Results on numerical accuracy for Ranger.

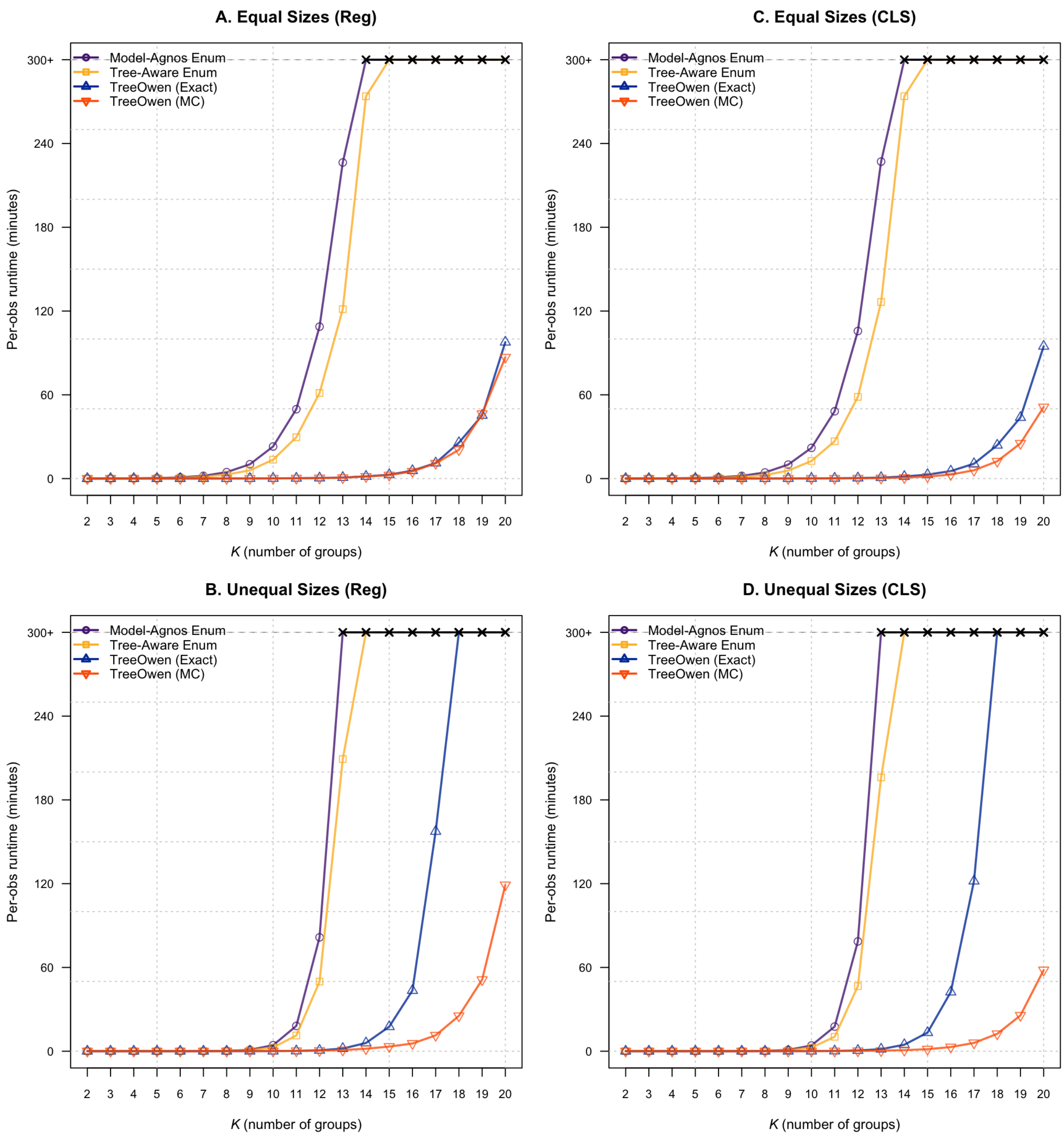


**Fig. S4.** Results on computational efficiency for Ranger.

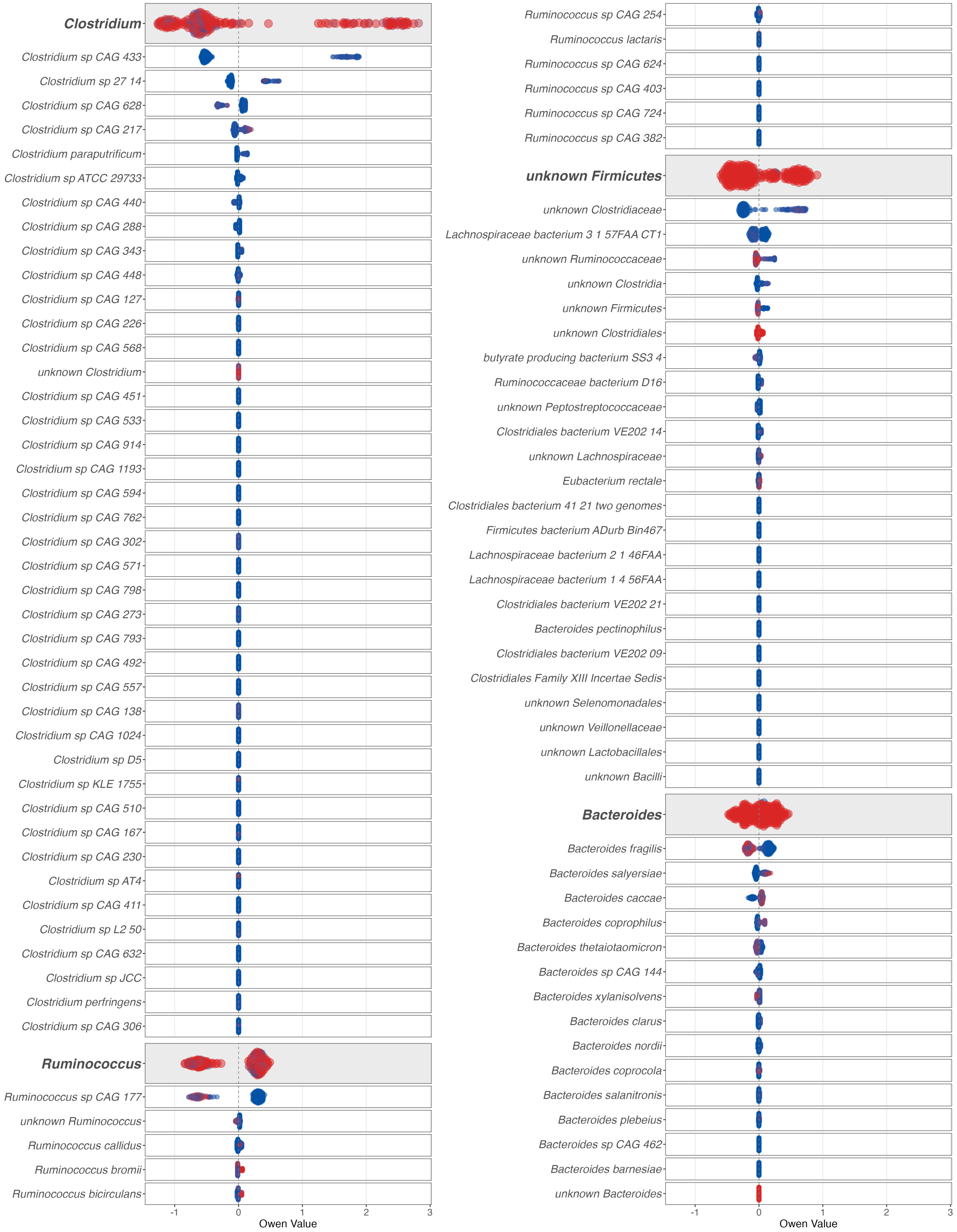


**Fig. S5.** First part of the full beeswarm plots for the real data application.

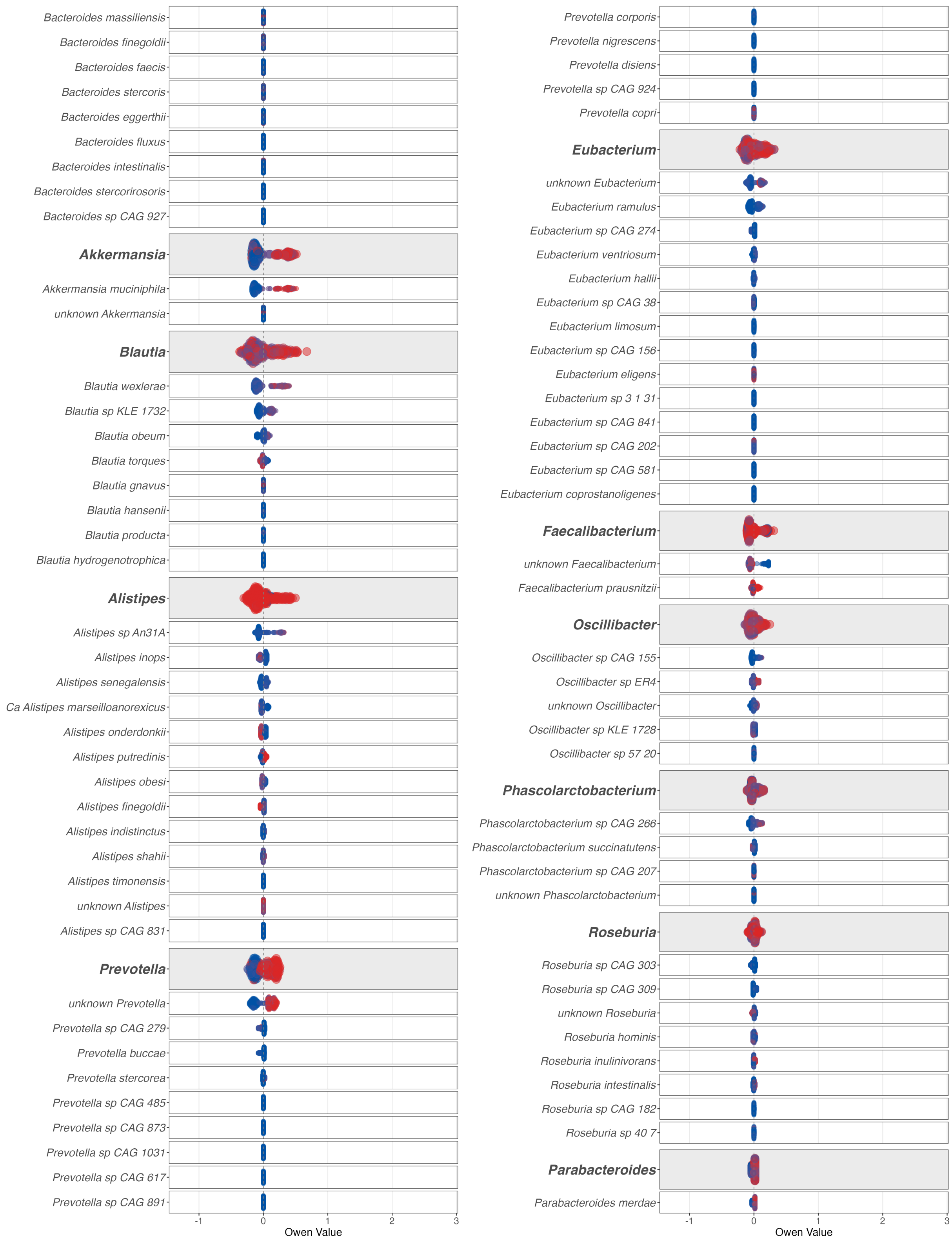


**Fig. S6.** Second part of the full beeswarm plots for the real data application.

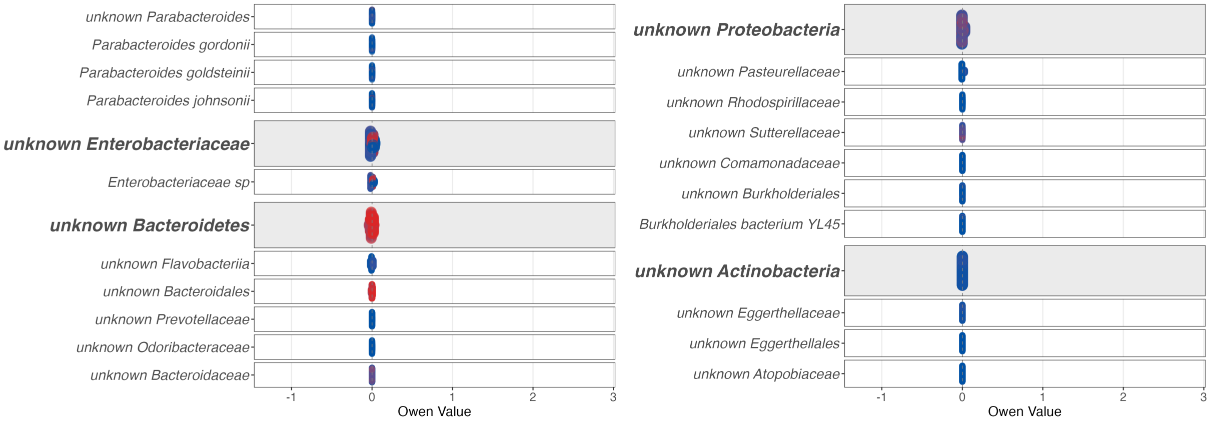


**Fig. S7.** Third part of the full beeswarm plots for the real data application.
